# HIDDEN APICOMPLEXAN PARASITE DIVERSITY LINKS CORAL AND PLANKTON MICROBIOMES ACROSS REEF SEASCAPES

**DOI:** 10.64898/2026.05.20.726672

**Authors:** Clément Leboine, Laura del Rio-Hortega, Nicolas Henry, Mathieu Zallio, Anthony M. Bonacolta, Caroline Belser, Jean-Marc Aury, Christian R Voolstra, Benjamin CC Hume, Alice Moussy, Clémentine Moulin, Emilie Boissin, Guillaume Bourdin, Guillaume Iwankow, Julie Poulain, Sarah Romac, Tara Pacific Consortium coordinators, Javier del Campo, Denis Allemand, Serge Planes, Maren Ziegler, Patrick Wincker, Quentin Carradec, Betina M. Porcel

**Affiliations:** Génomique Métabolique, Genoscope, Institut François Jacob, CEA, CNRS, Univ Evry, Université Paris-Saclay, Evry-Courcouronnes, France; Research Federation for the Study of Global Ocean Systems Ecology and Evolution, R2022/Tara Oceans GO-SEE, Paris, France; Department of Animal Ecology & Systematics, Justus Liebig University Giessen, 35392, Giessen, Germany; CNRS, Sorbonne Université, FR2424, ABiMS, Station Biologique de Roscoff, 29680, Roscoff, France; Department of Botany, University of British Columbia, Vancouver, BC V6T 1Z4, Canada; Department of Biology, University of Konstanz, Konstanz, Germany; Genoscope, Institut François Jacob, Commissariat à l’Energie Atomique (CEA), Université Paris-Saclay, 2 Rue Gaston Crémieux, 91057, Evry, France; Fondation Tara Océan, Base Tara, 8 rue de Prague, 75 012, Paris, France; PSL Research University: EPHE-UPVD-CNRS, USR CRIOBE, Laboratoire d’Excellence CORAIL, Université de Perpignan, Perpignan, France; School of Marine Sciences, University of Maine, Orono, Maine, United States of America; Sorbonne Université, CNRS, Station Biologique de Roscoff, AD2M, UMR, ECOMAP, Roscoff, France; Institut de Biologia Evolutiva (CSIC-Universitat Pompeu Fabra), Passeig Marítim de la Barceloneta 37-49, 08003, Barcelona, Catalonia, Spain; Centre Scientifique de Monaco, 8 Quai Antoine Ier, MC-98000, Principality of Monaco

**Keywords:** coral reefs, microbiome, parasite, Apicomplexa, metabarcoding

## Abstract

Parasitism is one of the most widespread trophic strategies in nature, though its diversity and ecological distribution in marine ecosystems remain poorly characterized. Apicomplexa are a major clade of obligate parasites best known for medically important taxa, yet their diversity and distribution in the ocean is still largely unresolved. Here, we used metabarcoding data from the Tara expeditions to investigate the diversity, distribution, and environmental drivers of Apicomplexa across coral reef ecosystems and adjacent oceanic habitats. By integrating samples spanning planktonic communities, coral tissues, and marine sediments across multiple oceanic regions, we substantially expand the known phylogenetic breadth of marine apicomplexans.

Although apicomplexans were generally low in relative abundance, they were widely distributed across marine environments. Community composition differed markedly among habitats. Corallicolid lineages were consistently associated with coral hosts, whereas planktonic samples harbored a greater diversity of apicomplexans, dominated by crustacean-associated gregarines. Sediments contained particularly high apicomplexan richness, including several poorly characterized groups. Capitalizing on the pan-Pacific transect of the expedition, we resolved biogeographic patterns in apicomplexan diversity across ocean basins: tropical regions showed the highest overall diversity, while polar environments contained distinct apicomplexan assemblages not detected in other ocean biomes.

Together, these results highlight the extensive and previously underappreciated diversity of marine Apicomplexa and demonstrate that integrating multiple marine biomes is essential for resolving the phylogenetic and ecological breadth of parasitism in the ocean.

## INTRODUCTION

Species interactions structure ecosystems, and parasitism is among the most prevalent and influential of these relationships. Parasitic protists of the phylum Apicomplexa are widespread across marine and terrestrial ecosystems [1, 2] and include major pathogens such as *Plasmodium falciparum* [3, 4] and *Toxoplasma gondii* [5]. These taxa have been extensively studied due to their medical and veterinary relevance [3–8], and their genomes have provided key insights into the evolution of parasitism [9–13]. However, they represent only a small fraction of total apicomplexan diversity, and the majority of lineages remain uncultured and poorly characterized in natural environments.

Environmental sequencing has revealed that apicomplexans are far more diverse and widespread than previously assumed, occurring across marine plankton, benthic sediments, coral reef systems, and soils [14–16]. These data suggest that parasitic eukaryotes may play underappreciated roles in regulating host populations and structuring microbial communities in natural ecosystems. However, most environmental lineages remain taxonomically unresolved, particularly among early-diverging marine groups such as gregarines and other poorly characterized apicomplexan clades [17, 18]. While their ecological roles and host associations remain incompletely resolved, available evidence suggests they may contribute to ecosystem functioning [14, 15, 19]. Still, this lack of resolution also limits our ability to assess how parasite diversity responds to environmental gradients and ecosystem structure at global scales.

Recent high-throughput environmental sequencing has further expanded the known diversity of the group through the discovery of novel apicomplexan and apicomplexan-related lineages, including coral-associated corallicolids, fish-infecting ichthyocolids, and marine Marosporida parasites infecting diverse invertebrate hosts [10, 15, 16, 20–30]. These findings indicate that substantial evolutionary diversity remains hidden within environmental datasets and that many lineages have yet to be linked to specific hosts or ecological functions. Despite this progress, global patterns of distribution and habitat specificity remain poorly constrained, particularly across contrasting marine ecosystems [16, 31–33].

Coral reefs and polar environments represent two ecologically and environmentally distinct marine systems that remain underrepresented in comparative surveys of microbial eukaryotes. Coral reefs are highly biodiverse and characterized by dense host–microbe hotspots [34–38], while polar oceans are shaped by low temperatures, strong seasonality, and distinct microbial community structures [31–33, 39, 40]. Characterizing apicomplexan diversity across these habitats provides a unique opportunity to explore ecological and biogeographic drivers shaping apicomplexan distribution at a global scale. Such contrasts are particularly powerful for disentangling whether parasite community structure is primarily shaped by host availability, environmental filtering, or geographic isolation.

Global oceanographic initiatives such as the Tara Pacific [41–43] and the Tara Polar [44] expeditions provide unprecedented access to microbial eukaryote diversity across coral reef-associated ecosystems, and to improve our understanding of apicomplexan diversity in polar oceans. Together, these coordinated sampling efforts span planktonic, benthic, coral-associated, and high-latitude environments, providing a comprehensive framework to investigate parasite diversity across broad spatial and ecological gradients in marine systems.

Here, we analyze 18S rRNA gene V9 metabarcoding datasets from Tara Oceans and Tara Pacific to characterize the diversity and environmental distribution of Apicomplexa across coral reef in the Pacific, encompassing sediment, coral and plankton biomes, as well as polar marine ecosystems. By integrating large-scale environmental sequencing with phylogenetic placement approaches, we aim to refine the evolutionary framework of marine apicomplexans and identify major ecological patterns shaping their distribution across contrasting oceanic biomes.

## MATERIALS AND METHODS

### Sample and Environmental Parameter Collection

Samples were collected using standardized protocols during the Tara Pacific expedition (2016–2018) [41], spanning corals, fish, plankton, and sediments (Supplementary Fig. 2). Coral microbiomes were sampled from three focal species (*Millepora platyphylla*, *Porites lobata*, *Pocillopora meandrina*) and additional broader colony sampling (hereafter CDIV) to capture wider host diversity. Taxonomic annotation of these was performed as described in [42, 45]. Seawater samples were collected across reef-associated gradients (coral surrounding waters (CSW), reef below surface (REEF SURFACE), and outside reef systems (NEAR ISLAND and OPEN OCEAN) and sequentially filtered into multiple size fractions (Supplementary Fig. 2). Sediment-associated microbiomes were additionally sampled near coral colonies. Environmental parameters are described in [42].

Samples from the Tara Oceans expedition (2009–2013) [44] were also included, spanning 210 stations across 20 biogeographic provinces. Sampling covered surface (SRF), deep chlorophyll maximum (DCM), mesopelagic zone (MESO), and mixed layer, (MIX) (collected when no clear DCM was detected) depths and size fractions from <3–5 µm to 180–2000 µm in (Supplementary Fig. 1B), as detailed in [46, 47].

### Metabarcoding Sequence Analysis

DNA extraction and library preparation protocols are detailed in [45] (Tara Pacific) and [48] (Tara Oceans). Amplicon Sequence Variants (ASV) abundance tables were generated using DADA2 [49] following published workflows available on Zenodo. Taxonomic was assigned using complementary approaches by IDTAXA [50] (confidence threshold 50%) and VSEARCH’s (best-hit matches with a sequence similarity > 80%) against the PR2 database (v5.0.0) [51].

### Metabarcoding data analysis

#### ASV filtering and Apicomplexa selection

The ASV abundance table was filtered prior to downstream analysis. First, ASVs with fewer than three reads per sample were removed from that sample to minimize potential background noise and PCR/artifacts. Subsequently, only sequences identified as eukaryotic using IDTAXA [50] were retained to exclude some prokaryotic and organellar 16S rRNA fragments co-amplified by the V9 18S rRNA primers.

Subsequently, the ASVs assigned to Apicomplexa using IDTAXA were extracted, together with ASVs displaying ambiguous classifications with IDTAXA but assigned to Apicomplexa by VSEARCH [52], as long as they fall within Apicomplexa-compatible IDTAXA higher-level clades (i.e. “unclassified_root”, “unclassified_Eukaryota”, “unclassified_TSAR”, and “unclassified_Alveolata”). Since both IDTAXA and VSEARCH used the PR2 reference database, the resulting annotations are directly comparable and share identical ranks. Taxonomic information was merged into a single annotation table for ASVs retained as Apicomplexa based on either method. IDTAXA confidence and VSEARCH sequence identity, together with the original assignment source, were kept for each ASV.

### Data normalization and abundance transformation

Relative abundance was computed (i) relative to the total eukaryotic community, and (ii) within Apicomplexa, providing a complementary viewpoint for global and Apicomplexa-focused interpretations.

We also applied a centered log-ratio (CLR) transformation [53] to make the data comparable and to reduce the impact of uneven sequencing [54]. A pseudocount of 1 was added prior to transformation, consistent with established approaches for handling zeros in compositional data [55]. For taxonomic abundance visualizations (e.g., genus-level barplots), ASVs were aggregated at the desired taxonomic rank prior to normalization.

### Alpha diversity and compositional overlap

UpSet plots were generated using the UpSetR package (v1.4.0) for visualization of compositional overlap among conditions (e.g., environment types, coral host genera).

Alpha diversity was computed in vegan (v2.6-4), with ASV richness (*specnumber()*) and Shannon diversity (*diversity()*). Group differences were tested using Wilcoxon tests with multiple testing Benjamini–Hochberg false discovery rate (FDR) correction [56, 57].

For spatial visualization, mean ± standard deviation (SD) Shannon values were calculated per island and environment type, and maps generated using ggplot2 (v3.5.2) with the “world” basemap (*map_data()*) in R [58].

### Beta diversity and multivariate community analyses

Samples with more than three Apicomplexa ASVs were retained for beta diversity analyses. Community structure was explored using principal component analysis (PCA) on centered log-ratio (CLR)-transformed abundances (*rda()* in vegan). Community differences were tested with PERMANOVA, alongside pairwise comparisons and analyses of dispersion heterogeneity (betadisper with permutation tests).

Beta diversity was calculated as Euclidean (Aitchison) distances on CLR-transformed data, with group differences tested by unpaired Wilcoxon tests and Benjamini–Hochberg correction [56, 57].

Environmental drivers of community variation were examined using redundancy analysis (RDA; *rda()* in vegan) on CLR-transformed abundances and standardized environmental variables. Variable selection was stratified by habitat type to maximize sample and variable coverage across planktonic and coral-associated datasets (Supplementary Fig. 4). Model significance was assessed using permutation-based ANOVA.

Optimal environmental subsets were identified with *bioenv()* (Spearman correlation). Relationships between community composition, environmental factors, and geographic distance were evaluated using Mantel and partial Mantel tests based on Euclidean distance matrices. Geographic distances were calculated as great-circle distances using the haversine formula implemented in distm() (geosphere v1.5-20). All analyses were performed in R [58] (v4.4.1), with figures generated using ggplot2 (v3.4.4).

### Phylogenetic placement

Environmental 18S rRNA (V9) sequences from Tara Oceans and Tara Pacific were phylogenetically placed onto a curated apicomplexan reference tree based on full-length and partial 18S rRNA sequences representing major lineages included in the PR2 framework [51]. This reference phylogeny was used to infer the evolutionary position of environmental ASVs within Apicomplexa.

V9 sequences were aligned to the reference multiple sequence using PaPaRa (v2.5) [59], and placed using phylogenetic placement methods [60]. Placement results were used to assign environmental sequences to major apicomplexan clades and to identify lineages lacking close reference representatives [61], and visualize in Anvi’o (v8) [62, 63].

In addition to confidently assigned ASVs, sequences with ambiguous or unresolved taxonomy were screened against the reference database to recover potential apicomplexan diversity missed by standard taxonomic assignment using BLASTn (v2.16.0) [64]. Only sequences consistent with apicomplexan affiliation based on similarity thresholds and placement confidence (715 ASVs; Supplementary Fig. 4A–B) were retained for downstream analyses. Final phylogenetic assignments were used to visualize diversity patterns across environmental samples with Krona (v2.8.1) [65].

### Comparison of Apicomplexa distribution and enrichment across Tara Oceans biomes and regions

An UpSet plot was generated using the UpSetR package (v1.4.0) in R, using 18S rRNA V9 metabarcoding data from the Tara Oceans expedition. For each ASV, presence at the biome level was defined as detection in at least one sample belonging to that biome.

ASV distributions across regions and biomes were visualized as bipartite networks using the ggraph (v2.2.1) and tidygraph (v1.3.1) packages. Apicomplexa enrichment (log-ratio) was modeled as a function of biome, depth, and size fraction using linear models. Linear mixed-effects models including region as a random intercept (lme4 v1.1-36) did not improve model fit (Akaike Information Criterion; AIC) and were not retained. Effects were tested by ANOVA in R, with pairwise comparisons based on estimated marginal means (emmeans v1.10.6) and Tukey adjustment. Interaction terms were included only when they significantly improved model fit.

Arctic and Southern Ocean samples were analyzed separately to assess regional differentiation. Linear models tested region, depth, and size fraction effects on enrichment. Post-hoc comparisons used estimated marginal means with Tukey-adjusted contrasts to compare regions within each depth.

## RESULTS

### Global distribution of apicomplexans across the marine seascape

Apicomplexans were consistently detected across Pacific tropical and subtropical reef environments, using 18S rRNA gene V9 metabarcoding datasets from the Tara expeditions [44, 66] Out of the 125,607 eukaryotic ASVs from the Tara Pacific survey (2.98 billion reads), 2,503 ASVs (2%) were classified as apicomplexans, spanning 114 taxonomic groups. These apicomplexan ASVs, identified across all Pacific biomes, accounted for nearly 10 million reads, representing 0.33% of the total eukaryotic sequence abundance (Fig. 1A). Among these ASVs, 381 sequences were shared between the two expeditions with 100% identity and coverage (Fig. 1C). This indicates that 61% of Tara Oceans ASVs (607 out of the 988 ASVs; Fig. 1A) and 85% of Tara Pacific ASVs (2,122 ASVs) were unique to each expedition (Fig. 1C), underscoring the strong added value of reef-associated and multi-biome sampling for capturing marine apicomplexan diversity.

**Figure 1.**
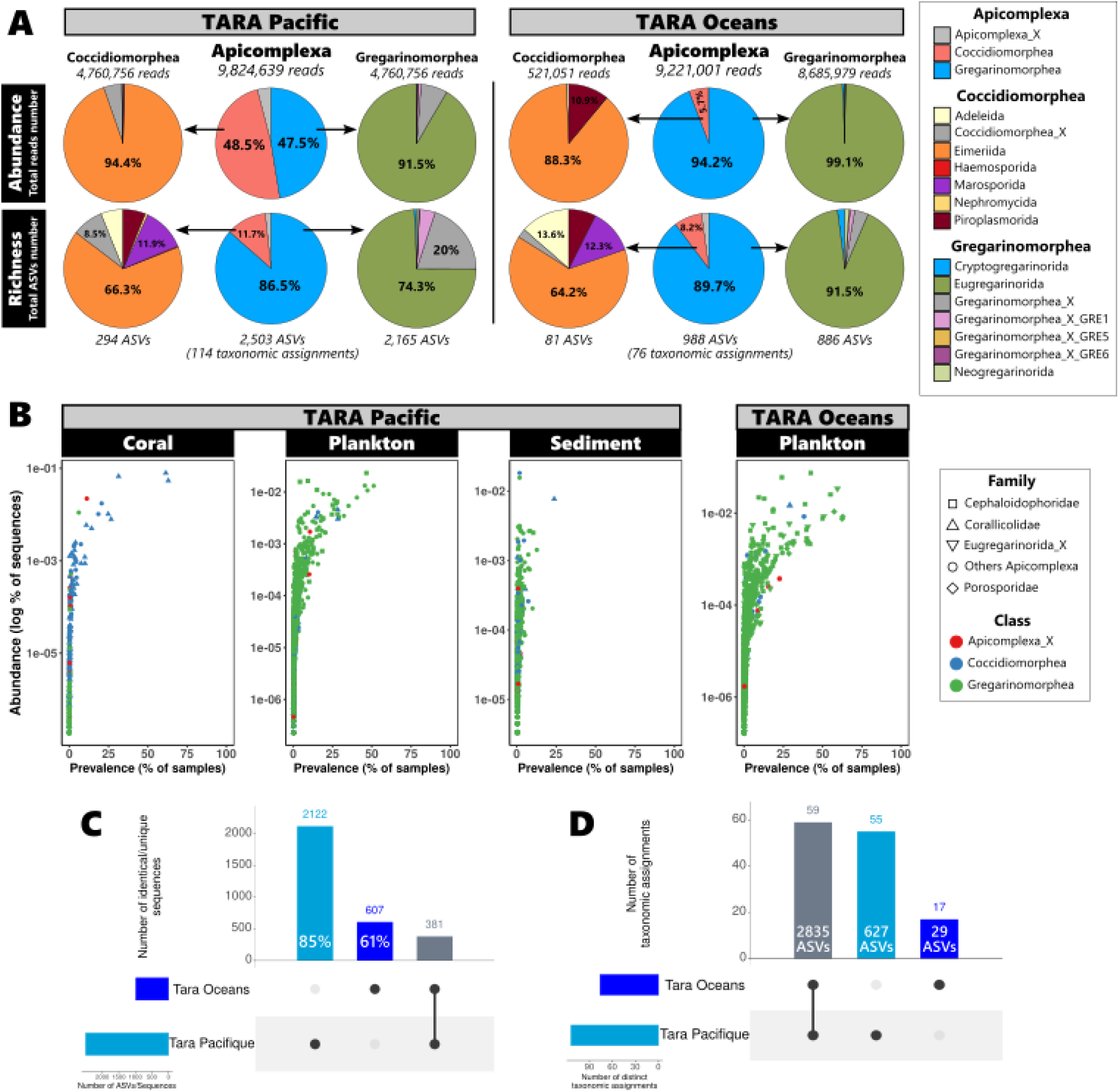
Global taxonomic distribution of the Apicomplexa phylum across oceanic environments. (A) Relative abundance (proportion of reads) and richness (proportion of ASVs) of Coccidiomorphea and Gregarinomorphea apicomplexans for Tara Pacific and Tara Oceans datasets. The total number of reads and the number of apicomplexan ASVs are indicated above and below each pie respectively. (B) Prevalence and abundance of each Apicomplexa ASV in corals and related anthozoan hosts, plankton and sediment samples from Tara Pacific and Tara Oceans expeditions. Colours indicate the Apicomplexa classes and point shapes represent the most abundant families, as defined by the PR2 taxonomy [51]. (C) Comparison of identical ASVs (100% identity and coverage) between Tara Pacific and Tara Oceans datasets. (D) Comparison Tara Pacific and Tara Oceans datasets at taxonomic level.

The Tara Oceans dataset contributed 98,358 eukaryotic ASVs (1.57 billion reads), expanding the survey to oceanic regions not covered by the Tara Pacific study. Among them, the 988 apicomplexan ASVs accounted for 76 taxonomic assignments and approximately 1% of the total eukaryotic ASVs (Fig. 1A).

The proportion of Apicomplexa-assigned ASVs varied across biomes. In non-reef oceanic environments (open ocean and near-island waters), relative abundance reached ∼1–1.5% of eukaryotic reads, but was lower in reef-associated habitats (reef surface and CSW), particularly in the 3–20 µm size fraction. In contrast, richness showed the opposite pattern: reef-associated environments harbored higher apicomplexan richness (∼2% of eukaryotic ASVs) than non-reef environments (∼1–1.5%) across most size fractions, except 0.2–3 µm (Supplementary Fig. 1A). In coral samples, relative abundance was below 0.5% for most coral genera, with slightly higher values in *Pocillopora* sp. and *Cyphastrea* sp., reaching up to 3% in *Acropora* sp.. Strikingly, apicomplexans were nearly absent from *Millepora* hydrozoans, and remained low in sediment samples (∼0.15% of eukaryotic reads and ∼0.5% of eukaryotic ASVs) (Supplementary Fig. 1A).

In contrast, apicomplexan richness was substantially higher than their relative abundance in corals, representing on average 2.7% of eukaryote ASVs and up to 6% in *Oulophyllia*, although based on a limited number of samples (Supplementary Fig. 1A).

Apicomplexan abundance varied strongly with both depth and size fraction (Supplementary Fig. 1). In seawater samples, the highest relative abundances were observed in the DCM and MES layers, accounting for up to 2-2.5% of total eukaryotic reads, whereas surface reef waters (SRF) consistently exhibited proportions below 1%. As for richness, apicomplexans represented 1-2% of eukaryotic ASVs, reaching 5% in the 0.8-5 µm size-fraction of the MES layer (Supplementary Fig. 1B).

Despite their relatively low overall abundance, apicomplexans were widespread in marine communities, occurring in 88% of Tara samples (6,655/7,568). Prevalence was highest in seawater and sediment (95% of samples; Supplementary Fig. 2). Among scleractinian corals, apicomplexans were detected in most genera (mean prevalence 87.8%), with the exception of the octocorals *Heliopora* (n = 6) and Alcyonacea (n = 1). The hydrozoan *Millepora* showed low prevalence (14.3% of samples; Supplementary Fig. 2).

### Taxonomic composition and dominance patterns of Apicomplexa

Coccidiomorphea and Gregarinomorphea, two major classes of the Apicomplexa phylum, were evenly represented in the Tara Pacific dataset, likely reflecting the high diversity of the sampled biomes (Fig. 1A). While Eimeriida, largely driven by the Corallicolidae family, was the most abundant group within Coccidiomorphea, the Sarcocystidae family constituted the second most ASV-rich family in Pacific environments, accounting for 8% of the total richness (Supplementary Table 1).

Conversely, the Gregarinomorphea largely dominated Tara Oceans seawater samples, while showing the highest richness across both datasets (86.5% of apicomplexan ASVs in Tara Pacific and 89.7% in Tara Oceans; Fig. 1A). The Cephaloidophoridae and Eugregarinorida_X families were consistently among the most abundant lineages of Gregarinomorphea across all seawater samples, despite the low diversity of crustacean-infecting gregarines (Fig. 1A and B; Supplementary Table 1). The Porosporidae family showed a balanced contribution to both abundance and richness. A similar pattern in Apicomplexa clade composition was observed in seawater samples from other ocean regions. (Supplementary Table 1).

Most apicomplexan ASVs exhibited low prevalence, occurring in fewer than 25% of samples. However, a small subset showed broader distributions (Fig. 1B). In coral samples, the most prevalent ASVs belonged to the family Corallicolidae, with two ASVs (ApiASV_1 and ApiASV_3) detected in approximately 70% of coral samples. Sediment communities were highly heterogeneous, with most ASVs displaying prevalence below 15%. Only one Corallicolidae ASV (ApiASV_123) departed from this pattern, reaching ∼25% prevalence exclusively in the sediment biome. In seawater samples, the most prevalent ASVs corresponding to Gregarinomorphea were assigned to the Cephaloidophoridae (ApiASV_6), Porosporidae (ApiASV_11 and ApiASV_15) and Eugregarinorida_X (ApiASV_12, ApiASV_19 and ApiASV_26) families, several of which reached prevalences between 50% and 75% (Fig. 1B).

### Expanding the known diversity of Apicomplexa

Phylogenetic placement of Tara Pacific and Tara Oceans ASVs onto a curated apicomplexan reference tree revealed extensive undescribed diversity across apicomplexan lineages.

Most Tara environmental sequences were confidently placed within the Apicomplexa reference phylogeny (Fig. 2), supported by predominantly low EDPL values and strong statistical support for a subset of placements (Supplementary Fig. 3 A-C). Few sequences were placed outside Apicomplexa, branching with outgroup references due to low placement support despite high sequence similarity to apicomplexan references (Fig. 2). This prevented confident taxonomic assignment and highlighted the limited characterization of marine apicomplexan diversity, particularly among deep-branching lineages.

**Figure 2.**
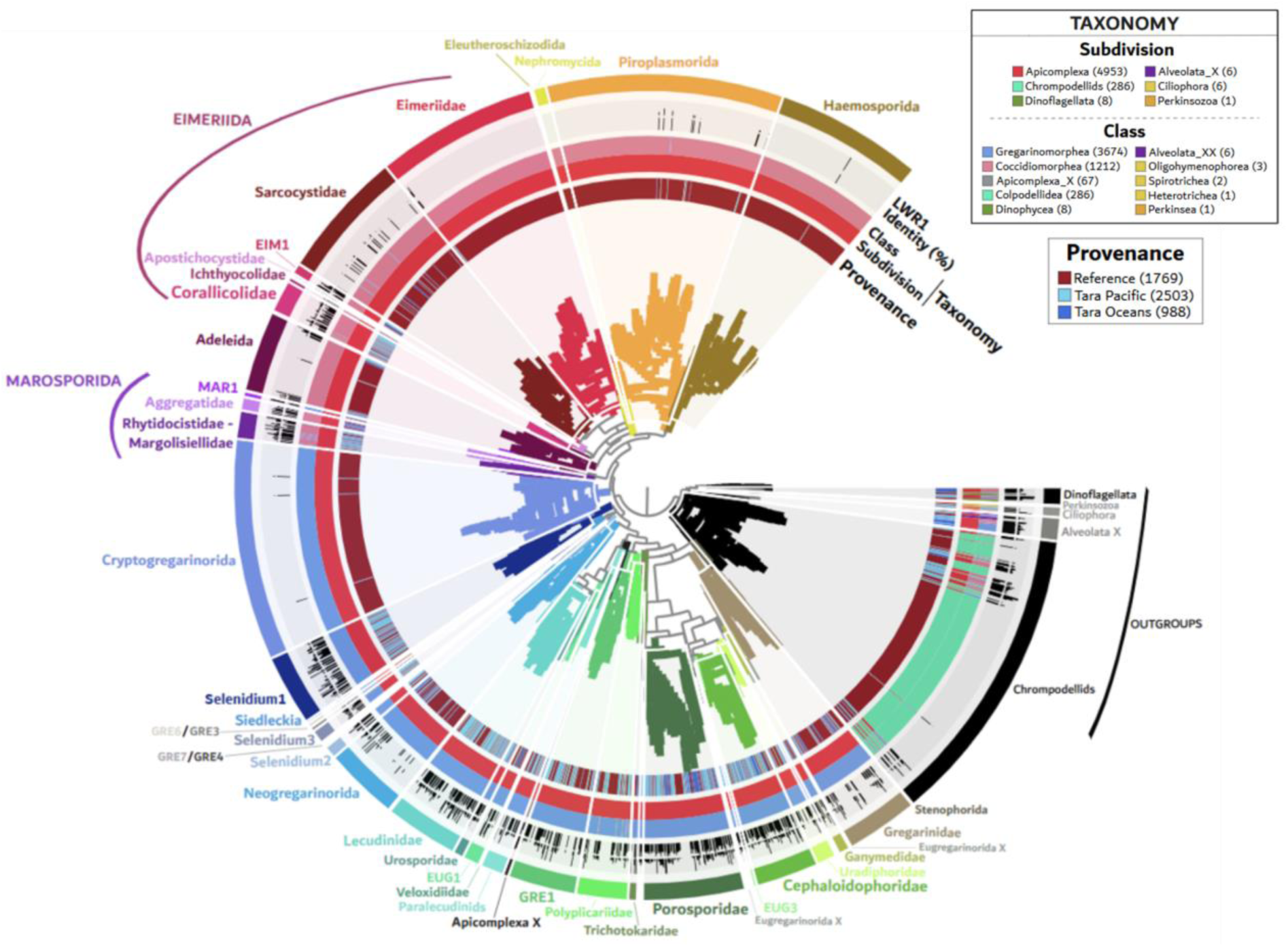
Phylogenetical placement of *Tara* apicomplexan ASVs onto a curated rRNA 18S reference tree. Environmental ASVs were placed on an apicomplexan reference phylogenetic tree, built using full-length and partial 18S rRNA sequences, and kindly provided by Dr. Javier del Campo. Phylogenetic placement was performed using a likelihood-based approach using EPA-ng, which computes the likelihood weight ratio (LWR) and the expected distance to the placement location (EDPL) for each query sequence, providing quantitative confidence measures for all feasible placements. The distribution of placement scores and the predominance of low EDPL values collectively indicate a satisfactory placement quality for most sequences globally (Supplementary Figure 3). The placement statistics were then summarized and filtered using GAPPA (v0.9.0) [61]. Visualization, annotation, and layer-based coloring of the final placement tree were carried out in Anvi’o (v8). From the outside in (i) The highest LWR1 for each placement (ranging from 0 to 1) representing the relative confidence of the best phylogenetic placement); (ii) percentage of identity between each ASV and the taxonomic assignment sequence (independent of the phylogenetic placement); (iii) High-level taxonomic assignation of 18S sequences, (iv) provenance, distinguishing reference sequences from ASVs originating from *Tara* Pacific or *Tara* Oceans. Major apicomplexan clades and families follow PR2 taxonomy [51]. Outgroups corresponding to non-apicomplexan lineages are displayed in black and grey.

The Tara environmental sequences were concentrated within several apicomplexan clades, notably the Marosporida class, the polyphyletic morphologically defined genus *Selenidium (sensus lato),* the Eugregarinorida order, and the Corallicolidae/Ichthyocolidae families (Fig. 2). These clades were enriched in environmental sequences relative to their number of described taxa, suggesting substantial higher diversity than currently recognized (Supplementary Table 2). This pattern was especially pronounced in Corallicolidae and Marosporida, with 2.3- and 1.5-fold enrichment, respectively (Fig. 2; Supplementary Table 2).

Environmental sequences were identified in other apicomplexan clades, albeit in lower proportions. For instance, 23% of the 18SV9 diversity observed in the Neogregarinorida order comes from Tara datasets, indicating the potential existence of marine representatives despite their predominantly terrestrial and freshwater insect hosts (Fig. 2 and Supplementary Table 2).

Marine apicomplexan diversity extends beyond what is captured by confidently assigned ASVs alone: some environmental sequences (n=29) could not be assigned to defined clades, forming basal or sister lineages to reference groups, reflecting limited reference coverage and likely undescribed phylogenetic diversity. Moreover, the phylogenetic placement of 715 ASVs retained as apicomplexan candidates adds new complexity to lineages such as Corallicolidae, Marosporida, and the gregarine clades. (Supplementary Fig. 4).

### Global community composition and diversity of apicomplexans across marine biomes

Apicomplexan community composition, diversity, and ASV turnover varied significantly across marine habitats and biomes sampled during the Tara expeditions, with the strongest compositional differences observed between coral- and seawater-associated communities (Fig. 3A, Supplementary Table 3). Comparisons involving sediment communities were also significant but explained less variance, likely due to higher within-group heterogeneity, as indicated by dispersion tests (Supplementary Table 3A and B). These differences are reflected in the taxonomic distribution of dominant genera across biomes (Fig. 3B).

**Figure 3.**
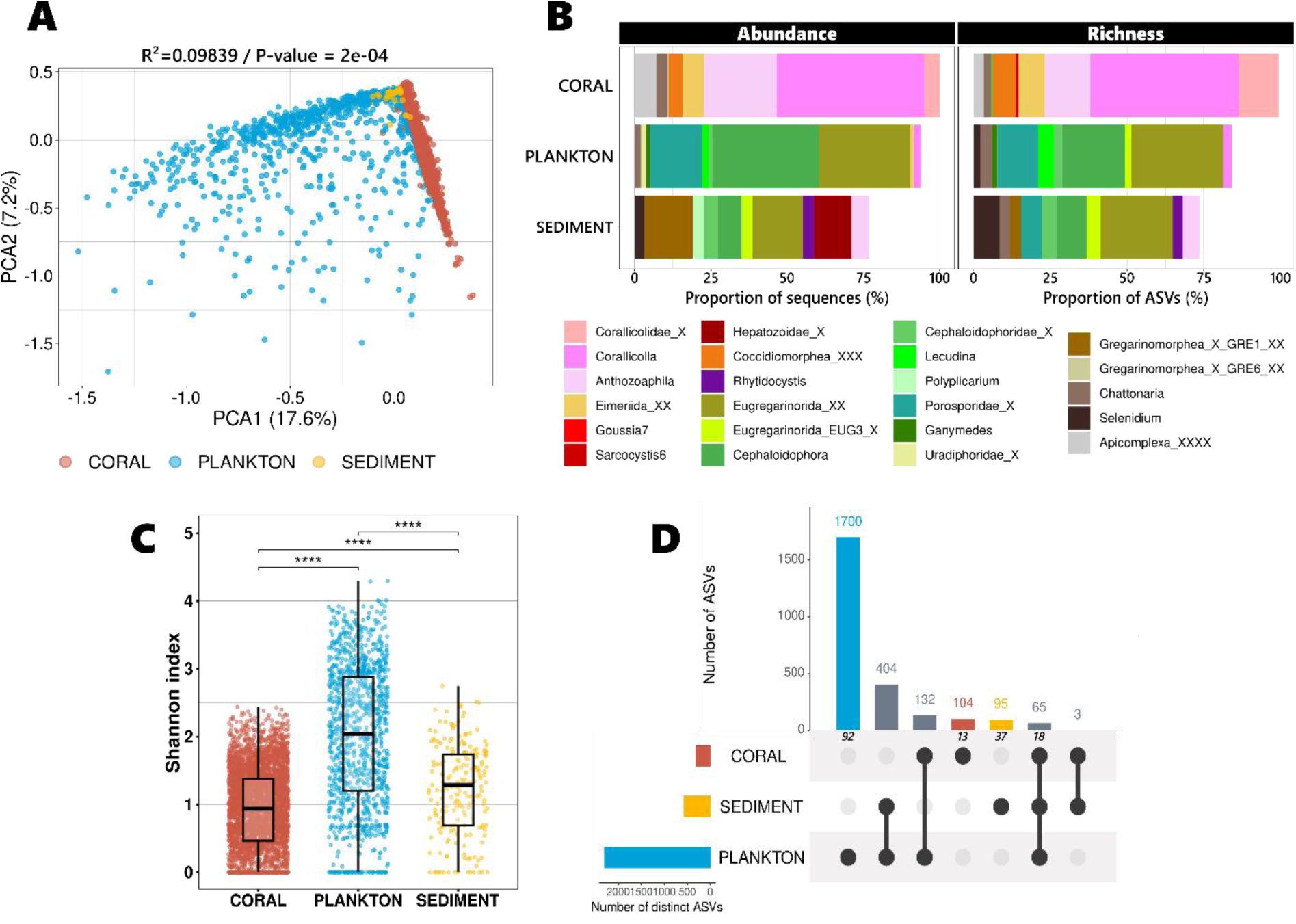
Community composition and diversity of apicomplexans across Pacific biomes and Tara expeditions. (A) PCA based on CLR-transformed abundance data showing differences in apicomplexan community composition between biomes. (B) Overall community composition for the 10 most abundant (left) and richest (right) Apicomplexa genera in the different Tara Pacific biomes. (C) Boxplot of Shannon diversity index across all samples with Apicomplexa showing significant differences between biomes (p ≤ 0.001). (D) UpSet plot showing the number of unique and shared ASVs among biomes. Italic numbers displayed below the upper bars indicate the corresponding number of different taxonomic assignments.

ASV-level analysis revealed strong contrasts in community structure among environments. Coral communities exhibited a high dominance structure, as the ten most abundant ASVs accounted for nearly 90% of the total abundance. Conversely, they represented only ∼40% of the total abundance in planktonic and sediment communities. Richness was more evenly distributed: the ten most diversified taxa represented ∼60% of total richness in coral samples, compared with ∼15% in plankton and sediment samples (Supplementary Fig. 5A–B).

Apicomplexan alpha diversity also differed significantly among biomes (Shannon index, Wilcoxon tests, p < 0.01). Planktonic communities exhibited the highest alpha-diversity, followed by sediment communities, whereas coral-associated communities displayed the lowest (Fig. 3C).

Shared and biome-specific ASVs were detected across Tara datasets. Only 14 ASVs were common to all four biomes, increasing to 65 ASVs when excluding fish samples (Fig. 3D). These shared ASVs were mainly gregarines (i.e., Cephaloidophoridae, Porosporidae, Eugregarinorida_X, and *Selenidium*), and, to a lesser extent, Corallicolidae. Overall, seawater samples harbored by far the largest number of specific ASVs (1,678 ASVs), mainly assigned to the families Lecudinidae (362 ASVs), Eugregarinorida_X (333 ASVs), Cephaloidophoridae (178 ASVs) and *Selenidium*-related lineages (Gregarinomorphea_XX, 313 ASVs) (Fig. 3D). Sediment samples contained fewer specific ASVs (94), notably including Rhytidocystidae (5 ASVs) and Gregarinomorphea_XX (31 ASVs; *Selenidium* and *Chattonaria*). As expected, coral samples were characterized by a high proportion of specific ASVs, among which 60% belonged to the family Corallicolidae (Fig. 3D).

### Worldwide distribution of seawater apicomplexans

In seawater samples, apicomplexan community composition was strongly structured between the open ocean, the vicinity of the islands and coastal samples within each size-fraction (Fig. 4A and Supplementary Table 3B). Similar patterns were observed across size fractions and depths in seawater samples collected beyond the Pacific (Fig. 4B).

**Figure 4.**
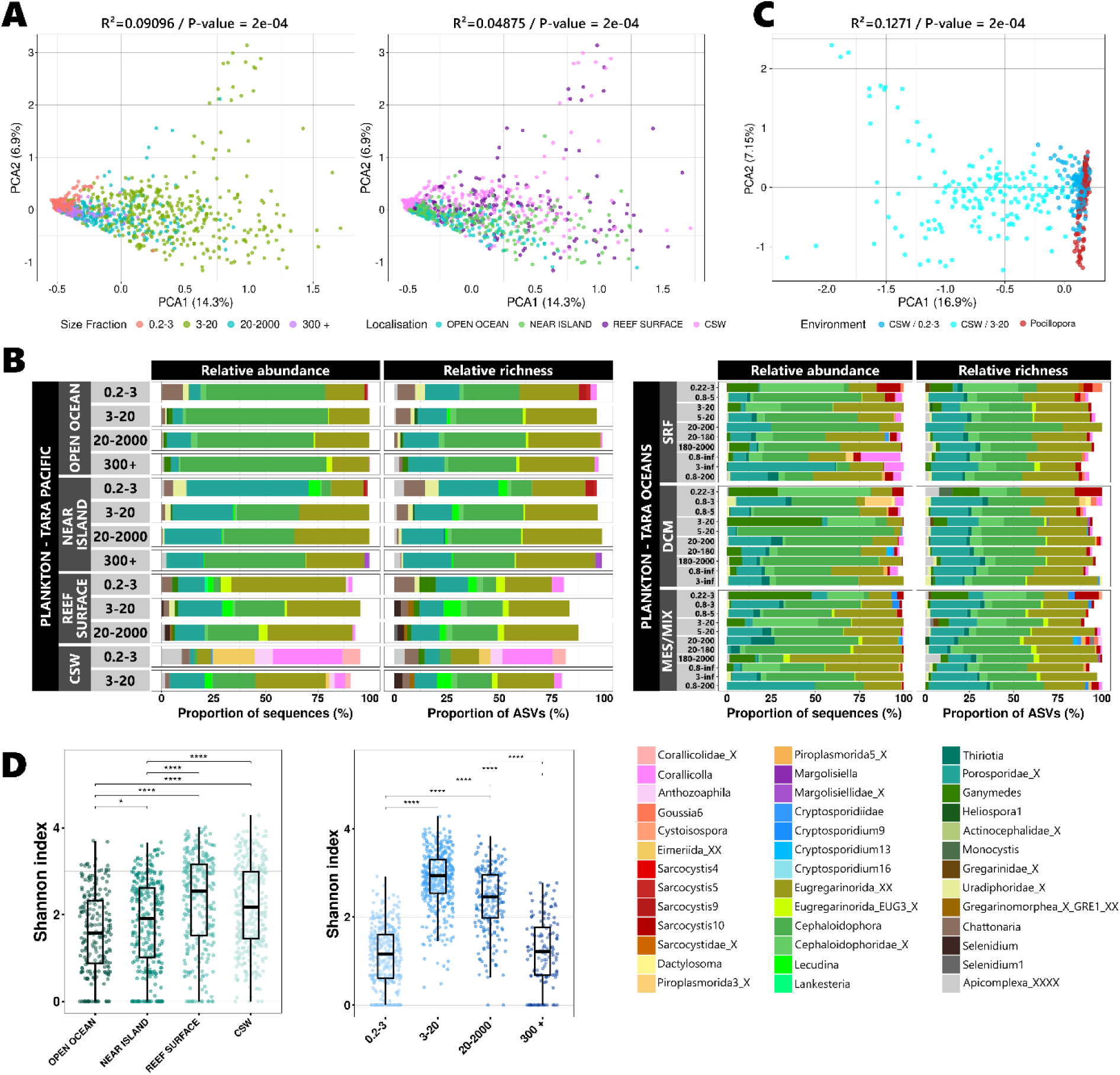
Community composition and diversity of Apicomplexa across seawater samples. (A) PCA based on CLR-transformed abundance data showing variation in Apicomplexa community composition across planktonic size fractions and oceanic environments in Tara Pacific samples. (B) Relative abundance and relative richness of the most abundant and richest Apicomplexa genera across planktonic size fractions in Tara Pacific (left) and Tara Oceans (right) samples. (C) PCA showing differences in Apicomplexa community composition between planktonic size fractions and coral reef surface waters (CSW). (D) Boxplots of Shannon diversity index values for planktonic communities grouped by oceanic environment and size fraction; significant differences (Wilcoxon test) are indicated by asterisks.

Along the distance to the coast, communities from coral-surrounding waters were the most distinct, differing significantly from both the ones near the islands and in open-ocean surface waters (Supplementary Table 3A). Along the size-fraction axis, the biggest differences were observed between the 0.2–3 µm fraction and the larger fractions (Fig. 4A and Supplementary Table 3A).

At the genus level, planktonic communities largely reflect the observed global patterns (Fig. 3B), with a dominance of gregarines like Eugregarinorida_XX, *Cephaloidophora* and Porosporidae_X across most size fractions and locations (Fig. 4B). These three genera also harbored the highest ASV richness, with a particularly high contribution of Eugregarinorida_XX (∼30%), and similar richness levels for Porosporidae and Cephaloidophoridae (∼20% each) (Fig. 4B). The main exception was the samples collected in the immediate vicinity of *Pocillopora* colonies, where corallicolids were more prominent, particularly within the 0.2–3 µm fraction. This resulted in a community structure that closely resembled coral-associated samples, possibly reflecting a sampling-related effect due to the presence of infected coral cells. (Fig. 4C and Supplementary Table 3A).

Apicomplexan communities became more diverse as sampling moved closer to reef habitats. For comparable size fractions, CSW and reef-surface samples consistently exhibited higher alpha diversity than near-island and open-ocean samples. The highest diversity was observed in the 3–20 µm fraction, harboring a high proportion of unique ASVs (Supplementary Fig. 6A), with similar diversity levels in CSW and reef-surface waters, whereas the 0.2–3 µm fraction was the least diverse across all locations (Fig. 4D). Diversity hotspots tended to cluster near coasts and islands across size fractions, particularly along the Asian margin, around Hawaii and in the Tropical Easter Pacific (Supplementary Fig. 7).

### Coral-associated apicomplexans are dominated by corallicolids, with important divergence across host genera

Coral-associated samples were overwhelmingly dominated by corallicolids, with higher apicomplexan abundance and richness across most coral hosts, often accompanied by ASVs with coarse or unresolved taxonomic assignments (e.g., Coccidiomorphea_XXX, Eimeriida_XX), which may correspond to additional, putative corallicolid lineages (Fig. 5A)

**Figure 5.**
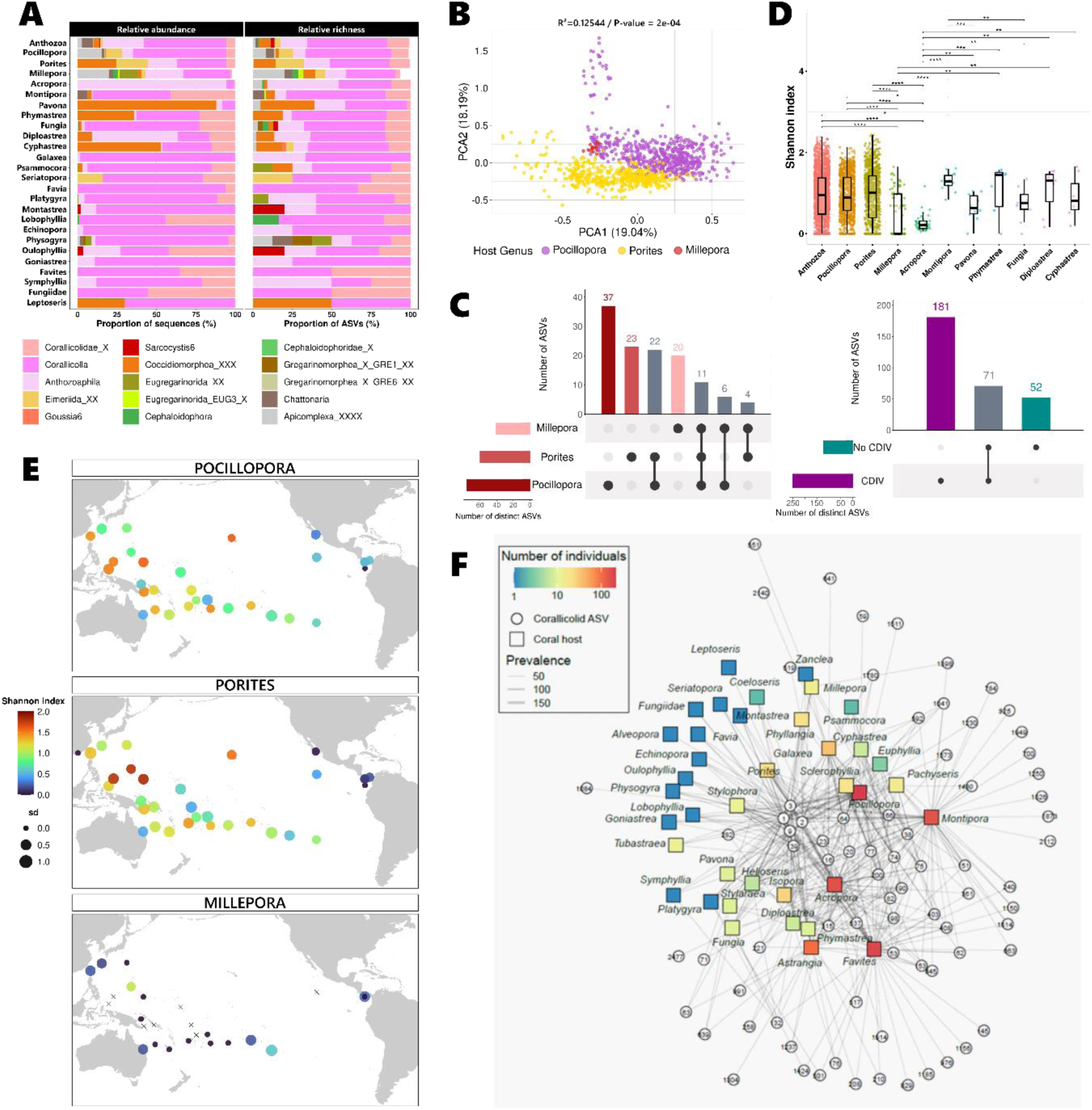
Community composition and diversity of Apicomplexa across coral host genera. (A) Relative abundance and relative richness of Apicomplexa taxa across coral host genera. (B) PCA based on CLR-transformed abundance data showing differences in Apicomplexa community composition among the coral genera *Pocillopora*, *Porites* and *Millepora* in Tara Pacific samples. (C) UpSet plots showing the number of shared and unique ASVs among coral genera (D). Boxplots of Shannon diversity index values for coral-associated communities grouped by host genus. Differences among groups were tested using unpaired Wilcoxon tests with Benjamini–Hochberg correction; significant differences are indicated by asterisks. (E) Geographic distribution of mean Shannon diversity index values per island for each coral genus; point size reflects the standard deviation among samples (F) Graph of corallicolid presence in different coral genera. Coral genera (squares) hosting corallicolid ASVs (circle) are connected with edges. Node positioning is optimised with the stress-minimization algorithm. The number of individual corals is indicated by the square colour. The edge width is proportional to the prevalence of the ASV in the coral genus. In (B) and (C) “Anthozoa” stands for “unidentified anthozoan corals”.

Still, community composition differed significantly between the two scleractinian hosts, *Pocillopora* and *Porites*, reflecting a consistent shift in taxonomic makeup rather than increased heterogeneity within either group (Fig. 5B, Supplementary Table 3A and B). As for the hydrozoan *Millepora*, the low proportion of Apicomplexa is associated with greater variability in community composition compared to the two scleractinian genera.

These differences were reflected in the distribution of dominant genera: at the ASV level, *Pocillopora*, (37) *Porites* (23) and *Millepora (20)* harboured genus-specific ASVs, many of which were assigned to Corallicolidae (Figure 5C). Broader coral sampling significantly enhanced apicomplexan diversity. CDIV samples revealed 181 additional ASVs undetected in target genera (Fig. 5C), highlighting the importance of host diversity in characterizing these communities.

*Pocillopora* and *Porites* exhibited broadly similar overall distributions of alpha-diversity (Figure 5D). Apicomplexans were ubiquitous for both host genera, exhibiting similar large-scale patterns and diversity maxima along the Indonesian and Australian coasts. However, island-scale patterns frequently diverged between genera (Fig. 5E), with intra-island diversity ranging from homogeneous to highly contrasting among sites (Supplementary Fig. 8A and B). For *Millepora*, diversity patterns were highly heterogeneous, with rare high-richness samples among mainly low-diversity ones (Fig. 5E). Only one island, Guam, displayed consistently higher *Millepora*-associated apicomplexan diversity with relatively homogeneous values across sites. In most other islands, *Millepora* either harbored very low and heterogeneous diversity or no detectable Apicomplexa at all (Supplementary Fig. 8C). For the other coral genera, scleractinians displayed marked differences in apicomplexan diversity, ranging from high richness in *Montipora* to relatively low diversity in *Acropora* (Fig. 5D).

To identify specific versus generalist corallicolid ASVs, a force-directed graph of the coral-corallicolid associations was built (Fig. 5F). This network revealed that the specificity of corallicolids to their hosts depends on the ASV taxa. A few generalist ASVs (1, 2, 3, 9, 16, 23, 39, etc.) were present in most of the sampled corals (center of the graph), while many rare variants were detected in only a few coral colonies and appeared to be host-specific (outer limits of the graph). Sampling more coral colonies of the same genus increases the likelihood of observing rare variants.

### Sediment communities as reservoirs of rare and habitat-specific apicomplexans

Sediment communities displayed a more even distribution of dominant taxa, as the ten most abundant genera together accounted for ∼75% of total abundance. As in seawater samples, Gregarinomorphea dominated the top-ranked sediment genera (∼55%), while a smaller contribution of Coccidiomorphea (∼20%) was also observed. Notably, several genera that were rare or absent from other environments were detected in sediments, including Gregarinomorphea_X_GRE1 among gregarines, and Hepatozoidae_X and *Rhytidocystis* (Marosporida) among coccidiomorphs (Fig. 3B and Fig. 6). In sediments, community composition was markedly heterogeneous between samples. Some were dominated by gregarines, others by corallicolids, and others by Marosporida, a newly established Apicomplexa clade of obligate parasites such as *Marosporidium daphniae*, *Enterocytospora*, and *Hepatospora*, mainly infecting aquatic crustaceans [29]. This sample-level heterogeneity makes sediment communities ideal for identifying rare or spatially restricted clades and their geographic hotspots. (Fig. 6). The clade Marosporida was detected in sediments from several islands in the South Pacific, including Niue (I09), Wallis and Futuna (I10) and Tuvalu (I12T), as well as from island Clipperton (I31), located near the American coast. The clade Gregarinomorphea_X_GRE1 was also found predominantly in sediments and reached high relative abundances in several islands, including Coiba (I02), Fiji (I18), Palau (I25) and Hong Kong (I27). Finally, this fine-scale analysis revealed the presence of Cryptogregarinorida, corresponding to vertebrate-infecting parasites such as *Cryptosporidium*, in several sediment samples. Previously undetected, this signal represents a significant fraction of apicomplexan abundance in specific samples, particularly at islands Rapa Nui (I04) and Gulf of California (I30) (Fig. 6).

**Fig. 6.**
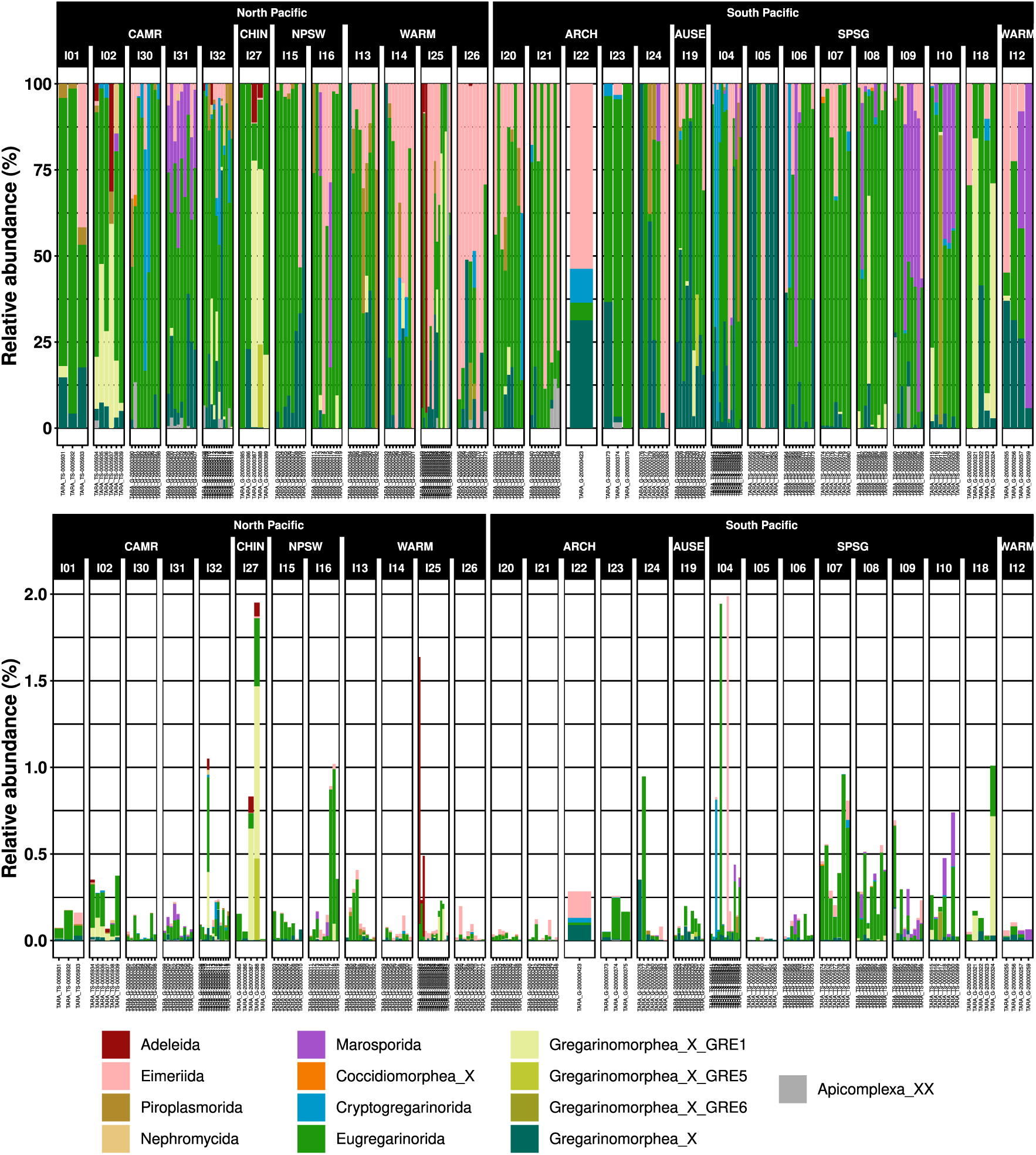
Spatial patterns of Apicomplexa diversity and community composition in sediment samples. (A) Relative abundance of Apicomplexa in sediment samples, colored by taxonomic order. The upper panel shows relative abundance within Apicomplexa, whereas the lower panel shows relative abundance relative to total eukaryotic communities. Longhurst biogeographical provinces are indicated by their code names: CAMR – Central American Coastal Province; CHIN – China Sea Coastal Province; NPSW – North Pacific Subtropical Gyre (West); WARM – Western Pacific Warm Pool Province; ARCH – Archipelagic Deep Basins Province (Indonesian region); AUSE – Australia–New Zealand Coastal Province (Eastern Australia); SPSG – South Pacific Subtropical Gyre. Islands are indicated by their code names, as described in [42].

### Influence of geographic and environmental factors on apicomplexan community structure

Apicomplexan community structure was primarily governed by island-scale spatial processes rather than large-scale oceanic structuring (Fig. 7A). This trend was particularly pronounced in *Acropora*, sampled only at two geographically distant islands (Ducie in Western Pacific and New Britain in Eastern Pacific). Comparable similarities within sites and between islands indicate that spatial structuring is driven primarily by island-level rather than site-level processes (Fig. 7A).

**Figure 7.**
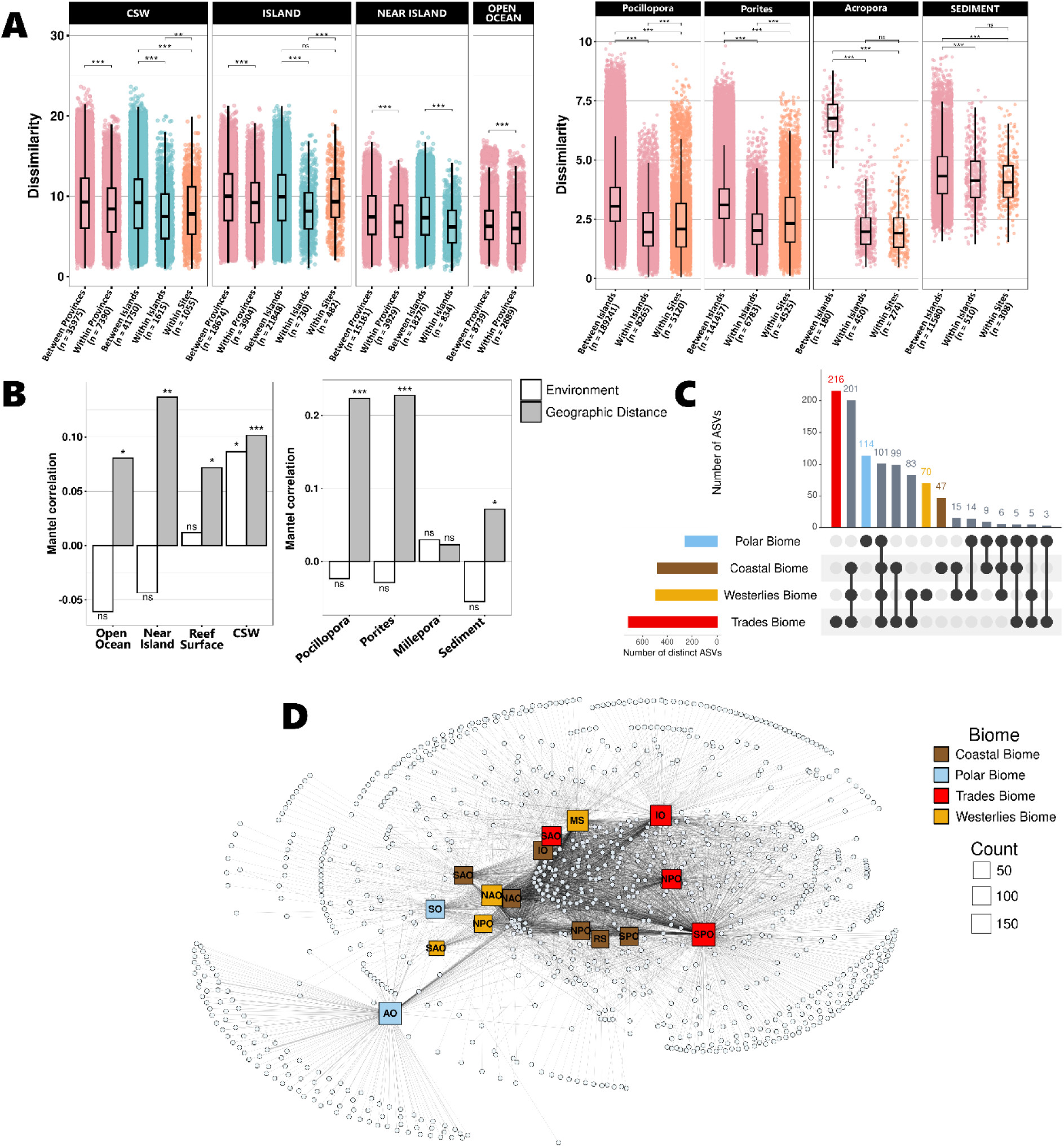
Patterns of beta diversity, Mantel correlations, and ASV connectivity across biomes. (A) Boxplots of pairwise dissimilarity of apicomplexan communities compared between provinces, within provinces, between islands, within islands and within sites at each island across oceanic biomes (left) and across coral host genera and sediment samples (right). Dissimilarity was calculated as the Euclidean distance of centered log-ratio (clr) transformed ASV data. Box plot horizontal bars indicate the median value, the box indicates the first and third QRs, and the whiskers indicate 1.5*IQR. N stands for the number of comparisons. (B) Mantel correlations between pairwise community dissimilarity and geographic distance or environmental distance for each oceanic biome (left) and coral and sediment samples (right). Bars show correlation coefficients, with significance indicated with stars. (C) UpSet plot showing the number of Apicomplexa ASVs that are unique to or shared among the Tara Oceans biomes (Trades, Westerlies, Coastal, Polar). An ASV is considered present in a biome if detected in at least one sample from that biome. (D) Bipartite network linking Apicomplexa ASVs to region × biome nodes. Circles represent ASVs and squares represent region–biome combinations (a region can include multiple biomes). Squares are colored by biome and labelled with region abbreviations (AO, Arctic Ocean; SO, Southern Ocean; SPO, South Pacific Ocean; NPO, North Pacific Ocean; IO, Indian Ocean; SAO, South Atlantic Ocean; NAO, North Atlantic Ocean; MS, Mediterranean Sea; RS, Red Sea). Square size is proportional to the number of samples available for each region–biome node. Edges indicate occurrence of an ASV in that region–biome and edge width reflect the number of samples in which the ASV is detected within that node.

While Mantel tests confirmed significant but weak distance-decay across all habitats, with stronger effects in near-island and coral surface waters, environmental filtering remained secondary and habitat-specific (Fig. 7B).

RDA marginal effects revealed habitat-specific drivers: oxygen and temperature in open oceans; phosphate and manganese near islands; and temperature, salinity, and phosphate in reef environments (Fig. 8A). Host-specific signatures also emerged for *Pocillopora* and *Porites*. However, the lack of strong multivariate environmental drivers across most habitats (Supp. Table 4) suggests that environmental influence is context-dependent and subordinate to the dominant spatial structuring observed at the island scale.

**Figure 8.**
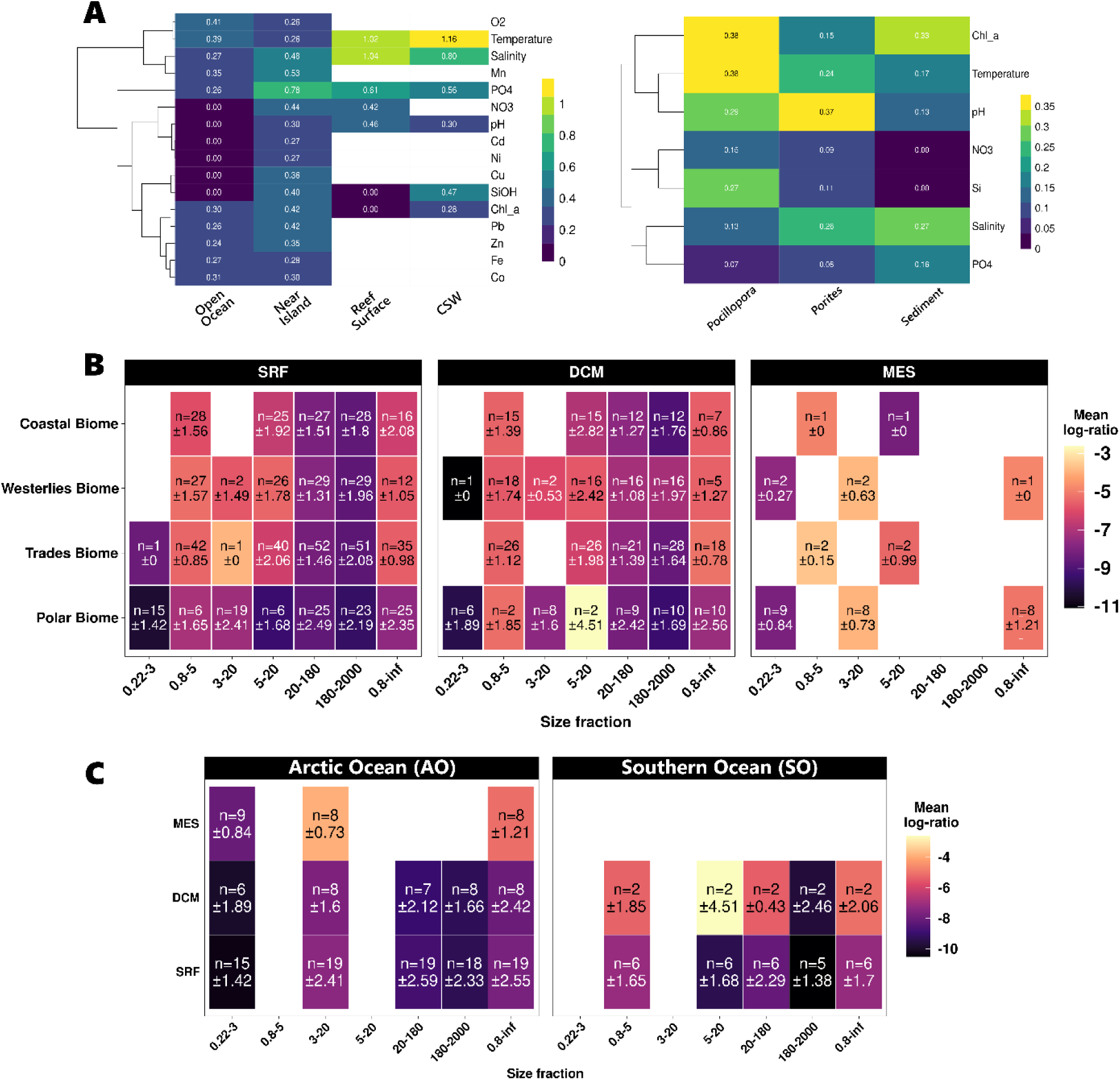
Environmental drivers of Apicomplexa community structure. (A) Heatmaps of the marginal variance explained (R²) by individual environmental variables in RDA models for oceanic biomes (top), coral and sediment samples (bottom); variables are ordered by hierarchical clustering based on euclidean distances. (B) Heatmap of Apicomplexa relative enrichment across size fractions, depth layers, and biomes. Enrichment is expressed as a log-ratio of Apicomplexa reads relative to non-Apicomplexa eukaryotic reads (higher values indicate higher relative enrichment of Apicomplexa). Tile color corresponds to the mean log-ratio for each combination. Each tile reports the number of samples (n) and the standard deviation (± SD). Depth layers: SRF (surface), DCM (deep chlorophyll maximum), MES (mesopelagic). Size fractions are indicated on the x-axis. (C) Same as (B) but restricted to the Polar biome, shown separately for the two polar regions (Arctic Ocean, AO; Southern Ocean, SO), with tiles reporting mean log-ratio, n, and ± SD for each depth × size-fraction combination.

### Apicomplexan distribution across oceanic provinces

Apicomplexan ASV distribution varied significantly across Longhurst provinces [67] (Fig. 7C). Biome-specific diversity peaked in the Trades (tropical open ocean), followed by the Polar biome, while the Westerlies and Coastal biomes showed fewer unique ASV counts. Although low- and mid-latitudes shared many ASVs, a core group of apicomplexans spanned all four biomes. Bipartite network analysis further resolved this spatial structuring (Fig. 7D), identifying the Arctic as a distinct module characterized by high endemism. This highlights that, although some lineages span broad ecological ranges, polar communities show strong regional diversity.

Apicomplexa enrichment varied significantly across biomes, depths, and size fractions (GLM; all p < 2.2e-16; Fig. 8B). Polar biomes showed lower enrichment than lower-latitude regions (p < 0.001), whereas mesopelagic layers and intermediate size fractions (0.8–20 µm) exhibited the highest values. Within polar regions, a significant Region × Depth interaction (p = 0.00089) revealed that the Arctic was more enriched than the Southern Ocean at the DCM (p = 0.0008) but comparable at the surface (Fig. 8C). Yet, insufficient Southern Ocean samples precluded mesopelagic comparisons. Altogether, polar community differences are primarily driven by vertical gradients (depth) rather than size-fractions, with the Arctic–Antarctic divergence peaking in the DCM.

## DISCUSSION

### Global Resolution of the Marine Apicomplexan Landscape

Our study provides a comprehensive global resolution of marine apicomplexan diversity, demonstrating that marine Apicomplexa represent a significant yet historically underappreciated component of global ocean diversity. While previous surveys highlighted the dominance of *Syndiniales* in marine habitats [68–70], our results reveal that apicomplexans are ubiquitous, exhibiting substantial abundance and evolutionary complexity across pelagic, marine benthic, and reef habitats, and are among the dominant parasite groups. By resolving these global distribution patterns, we fundamentally challenge the traditional view of Apicomplexa as primarily vertebrate-associated parasites [71, 72]. These findings underscore the value of global genomics initiatives like Tara missions in uncovering ‘cryptic’ parasite diversity and fill a vital gap in our understanding of the ocean’s ecological landscape.

### The Apicomplexan Paradox: Global Sparsity vs. Local Dominance

Marine apicomplexans exhibit a striking ecological paradox: they are globally sparse in terms of total eukaryotic richness yet geographically ubiquitous across all examined habitats [14–16]. Our pole-to-pole analysis confirms that while these parasites contribute minimally to global richness, they achieve localized dominance in specific niches, particularly within coral holobionts. This “patchy” distribution likely reflects a « boom-or-bust » parasitic lifestyle, where detection is decoupled from global abundance and governed by host availability and life-cycle synchronization [73, 74].

### Specialization and the Abundance-Diversity Decoupling

We observed a fundamental divergence in ecological strategies among gregarines. The crustacean-parasitic members of Cephaloidophoridae, which accounted for half of the apicomplexan in our study, displayed a sharp decoupling of abundance and diversity, characterized by high local dominance despite low ASV richness. This is a hallmark of ecological specialists adapted to constrained host requirements or microhabitat conditions [75], paralleling the strategy of *Syndiniales* Group II, whose global distribution and community structure is tightly constrained by sea surface temperature and nutrient-driven host availability [15, 69].

Conversely, members of Porosporidae exhibited a balanced abundance–diversity relationship, suggesting moderate ecological versatility or broader host ranges, similar to cosmopolitan generalists like the ciliate *Strombidium* spp., whose distribution has weak associations with temperature and nutrient gradients [15].

### Coral-Associated Dynamics: Host Specificity and Symbiosis

Coral diversity is mainly driven by corallicolids in our study. Corallicolids exhibit broad host and habitat distributions across anthozoan lineages, with evidence of host switching and variable host specificity [10, 23, 24, 76]. Our data expands the existing knowledge of potential corallicolid host specificity—including the first record of corallicolids in the hydrocoral *Millepora*—consistent with a flexible, generalist ecological strategy, suggesting a long-term symbiotic or low-virulence parasitic lifestyle. Previous studies have reported high corallicolid prevalence in healthy *Acropora* and the absence of consistent pathogenic symptoms, suggesting a commensal rather than classical parasitic role in this host [30]. This hypothesis might suggest that apicomplexans are not direct drivers of bleaching. Instead, their increased relative abundance in stressed corals likely reflects holobiont disturbance rather than a causal role [77]. However, this interpretation is not exclusive: while corallicolids were not shown to correlate with heat-stress–induced necrosis in octocorals [26], wno causal link to bleaching could be demonstrated in [77]. Overall, the evidence supports a flexible, non-binary view in which their ecological role cannot yet be confirmed or excluded and may depend on host state and environmental stress.

### Environmental Filtering vs. Spatial Structuring, Biogeographic Partitioning and the Island-level Effect

Apicomplexan community structure is primarily governed by geographic distance, with environmental parameters exerting a secondary, context-dependent influence. While individual variables explain a small fraction of total variance, we identified distinct “environmental signatures” for specific habitats. Composition was primarily associated with water-mass properties (oxygen and temperature) in seawater, as observed for other parasitic protists, which jointly regulate host availability and drive rapid community turnover [78]. Conversely, reef-associated and near-island communities were structured by productivity-related factors, specifically phosphate and manganese availability. Parasite communities in reef-associated plankton are shaped by productivity, with phosphate modulating host availability and manganese influencing primary production and holobiont physiology, thereby indirectly structuring parasite dynamics [78–81].

The community composition in sediments was highly heterogeneous, with low-prevalence taxa including rare clades consistently detected. A single Corallicolidae ASV exclusively present in this biome deviated from this pattern. However, this likely reflects a “sink” effect—the accumulation of coral-derived eDNA from mucus or detritus—rather than active sediment-adapted populations. Such sedimentary environmental DNA (eDNA) signals are well documented for reef-associated taxa and integrate biological inputs over broader spatial and temporal scales than water-column samples [82, 83].

Tropical systems exhibit higher diversity, driven by long-term climatic stability and host diversification [16, 84], whereas polar biomes are dominated by more specialized, seasonally structured lineages [31, 85]. Our results suggest that relative abundance patterns across biomes are driven more by depth and ocean basin than by a simple polar–tropical dichotomy. Strong island-level structuring across biomes reflects dispersal constraints, with islands acting as discrete biogeographic units shaped by isolation and ocean currents rather than cosmopolitan dispersal.

### The Frontier of Unresolved Diversity

By integrating Tara Oceans Arctic data with Pacific coral metabarcoding, we substantially extend the foundational work of del Campo et al. (2019). From marine sediments full of cryptic symbionts to insect guts containing specialized parasites, Apicomplexa emerge as far more ecologically and evolutionarily diverse than previously recognized [16, 27, 86]. Our survey enriched previously underrepresented branches, like the EM1 (Goussia-like) and Piroplasmorida-like clades. Both are obligate vertebrate parasites, and their detection in the water column likely reflects eDNA from transmission stages, host material, or unsampled infections, suggesting dispersal across oceanic regions beyond directly observed infections [15, 87, 88]. Given the limited resolution of short 18S markers, these ASVs may represent related, poorly characterized marine lineages. We therefore conservatively assign them as EM1 (Goussia-like) and Piroplasmorida-like apicomplexans, reflecting a broader parasitic component of marine microbial eukaryotes rather than active host–parasite interactions.

Our most significant finding is the dominance of unclassified gregarines (Eugregarinorida_X), which represent the largest fraction of both richness and abundance in this study. The success of these taxonomically unresolved lineages suggests they comprise vast cryptic species complexes with diverse, specialized roles within the coral holobiont. These results emphasize that the current marine apicomplexan tree of life is profoundly incomplete. Resolving this “hidden” diversity will require a shift toward integrative taxonomic approaches that combine large-scale metabarcoding with host-specific ecological data.

### Concluding Remarks

By moving beyond host-targeted studies, this large-scale metabarcoding approach has revealed a hidden world of marine apicomplexan diversity. While these high-throughput analyses cannot provide direct observations of host-parasite interactions, they highlight critical environmental “hotspots” and novel lineages—such as *Rhytidocystis* and the recently identified Marosporida—for future targeted research. Resolving this taxonomic frontier will require an integrative approach that bridges environmental genomics with host-specific ecology to fully map the functional landscape of the ocean’s parasites.

### Tara Pacific consortium coordinators

Sylvain Agostini (orcid.org/0000-0001-9040-9296) Shimoda Marine Research Center, University of Tsukuba, 5-10-1, Shimoda, Shizuoka, Japan

Denis Allemand (orcid.org/0000-0002-3089-4290) Centre Scientifique de Monaco, 8 Quai Antoine Ier, MC-98000, Principality of Monaco

Emilie Boissin (orcid.org/0000-0002-4110-790X) PSL Research University: EPHE-UPVD-CNRS, USR 3278 CRIOBE, Laboratoire d’Excellence CORAIL, Université de Perpignan, 52 Avenue Paul Alduy, 66860 Perpignan Cedex, France

Chris Bowler (orcid.org/0000-0003-3835-6187) Institut de Biologie de l’Ecole Normale Supérieure (IBENS), Ecole normale supérieure, CNRS, INSERM, Université PSL, 75005 Paris, France

Colomban de Vargas (orcid.org/0000-0002-6476-6019) Sorbonne Université, CNRS, Station Biologique de Roscoff, AD2M, UMR 7144, ECOMAP 29680 Roscoff, France & Research Federation for the study of Global Ocean Systems Ecology and Evolution, FR2022/ Tara Oceans-GOSEE, 3 rue Michel-Ange, 75016 Paris, France

Eric Douville (orcid.org/0000-0002-6673-1768) Laboratoire des Sciences du Climat et de l’Environnement, LSCE/IPSL, CEA-CNRS-UVSQ, Université Paris-Saclay, F-91191 Gif-sur-Yvette, France

Michel Flores (orcid.org/0000-0003-3609-286X) Weizmann Institute of Science, Department of Earth and Planetary Sciences, 76100 Rehovot, Israel

Didier Forcioli (orcid.org/0000-0002-5505-0932) Université Côte d’Azur, CNRS, INSERM, IRCAN, Medical School, Nice, France and Department of Medical Genetics, CHU of Nice, France

Paola Furla (orcid.org/0000-0001-9899-942X) Université Côte d’Azur, CNRS, INSERM, IRCAN, Medical School, Nice, France and Department of Medical Genetics, CHU of Nice, France

Pierre Galand (orcid.org/0000-0002-2238-3247) Sorbonne Université, CNRS, Laboratoire d’Ecogéochimie des Environnements Benthiques (LECOB), Observatoire Océanologique de Banyuls, 66650 Banyuls sur mer, France

Eric Gilson (orcid.org/0000-0001-5738-6723) Université Côte d’Azur, CNRS, Inserm, IRCAN, France

Fabien Lombard (orcid.org/0000-0002-8626-8782) Sorbonne Université, Institut de la Mer de Villefranche sur mer, Laboratoire d’Océanographie de Villefranche, F-06230 Villefranche-sur-Mer, France

David Arturo Paz-Garcia (orcid.org/0000-0002-1228-5221) Centro de Investigaciones Biológicas del Noroeste (CIBNOR), Laboratorio de Genética para la Conservación, Av. IPN 195, 23096, La Paz, BCS, México

Stéphane Pesant (orcid.org/0000-0002-4936-5209) European Molecular Biology Laboratory, European Bioinformatics Institute, Wellcome Genome Campus, Hinxton, Cambridge CB10 1SD, UK

Serge Planes (orcid.org/0000-0002-5689-5371) PSL Research University: EPHE-UPVD-CNRS, USR 3278 CRIOBE, Laboratoire d’Excellence CORAIL, Université de Perpignan, 52 Avenue Paul Alduy, 66860 Perpignan Cedex, France

Stéphanie Reynaud (orcid.org/0000-0001-9975-6075) Centre Scientifique de Monaco, 8 Quai Antoine Ier, MC-98000, Principality of Monaco

Shinichi Sunagawa (orcid.org/0000-0003-3065-0314) Department of Biology, Institute of Microbiology and Swiss Institute of Bioinformatics, Vladimir-Prelog-Weg 4, ETH Zürich, CH-8093 Zürich, Switzerland

Olivier Thomas (orcid.org/0000-0002-5708-1409) Marine Biodiscovery Laboratory, School of Chemistry and Ryan Institute, National University of Ireland, Galway, Ireland

Romain Troublé (ORCID not-available) Fondation Tara Océan, Base Tara, 8 rue de Prague, 75 012 Paris, France

Rebecca Vega Thurber (orcid.org/0000-0003-3516-2061) Oregon State University, Department of Microbiology, 220 Nash Hall, 97331Corvallis OR USA

Christian R. Voolstra (orcid.org/0000-0003-4555-3795) Department of Biology, University of Konstanz, 78457 Konstanz, Germany

Patrick Wincker (orcid.org/0000-0001-7562-3454) Génomique Métabolique, Genoscope, Institut François Jacob, CEA, CNRS, Univ Evry, Université Paris-Saclay, 91057 Evry, France

Maren Ziegler (orcid.org/0000-0003-2237-9261) Department of Animal Ecology & Systematics, Justus Liebig University Giessen, 35392 Giessen, Germany

Didier Zoccola (orcid.org/0000-0002-1524-8098) Centre Scientifique de Monaco, 8 Quai Antoine Ier, MC-98000, Principality of Monaco

## Acknowledgments

The *Tara* Pacific expedition would not have been possible without the participation and commitment of over 200 scientists, the Tara Ocean Foundation and its sailors, artists, staff and partners, the R/V Tara crew, and the Tara Pacific Expedition Participants (10.5281/zenodo.3777760). We are keen to thank the commitment of the following institutions for their financial and scientific support that made this unique Tara Pacific Expedition possible: le Centre national de la recherche scientifique (CNRS), l’Université Paris Sciences & Lettres (PSL), le Centre Scientifique de Monaco (CSM), l’École Pratique des Hautes Études (EPHE), le Centre National de Séquençage - Genoscope, le Commissariat à l’énergie atomique et aux énergies alternatives (CEA), Inserm, l’Université Côte d’Azur, l’Agence Nationale de la Recherche (ANR), Agnès Troublé said agnès b., UNESCO-IOC, the Veolia Foundation, the Prince Albert II de Monaco Foundation, Région Bretagne, Billerudkorsnas, AmerisourceBergen Company, Lorient Agglomération, Oceans by Disney, L’Oréal, Biotherm, France Collectivités, Fonds Français pour l’Environnement Mondial (FFEM), Etienne Bourgois, and the Tara Ocean Foundation teams. Tara Pacific would not exist without the continuous support of the participating institutes. The authors also particularly thank Serge Planes, Denis Allemand, and the Tara Pacific consortium. This project was funded by France Génomique (ANR-10-INBS-09). We also acknowledge C. Scarpelli and his team for support in high-performance computing, and the Genoscope Sequencing Team (https://zenodo.org/records/14611490) for the generation of the sequencing data.

## Supplementary Figures

**Supplementary Fig. 1.**
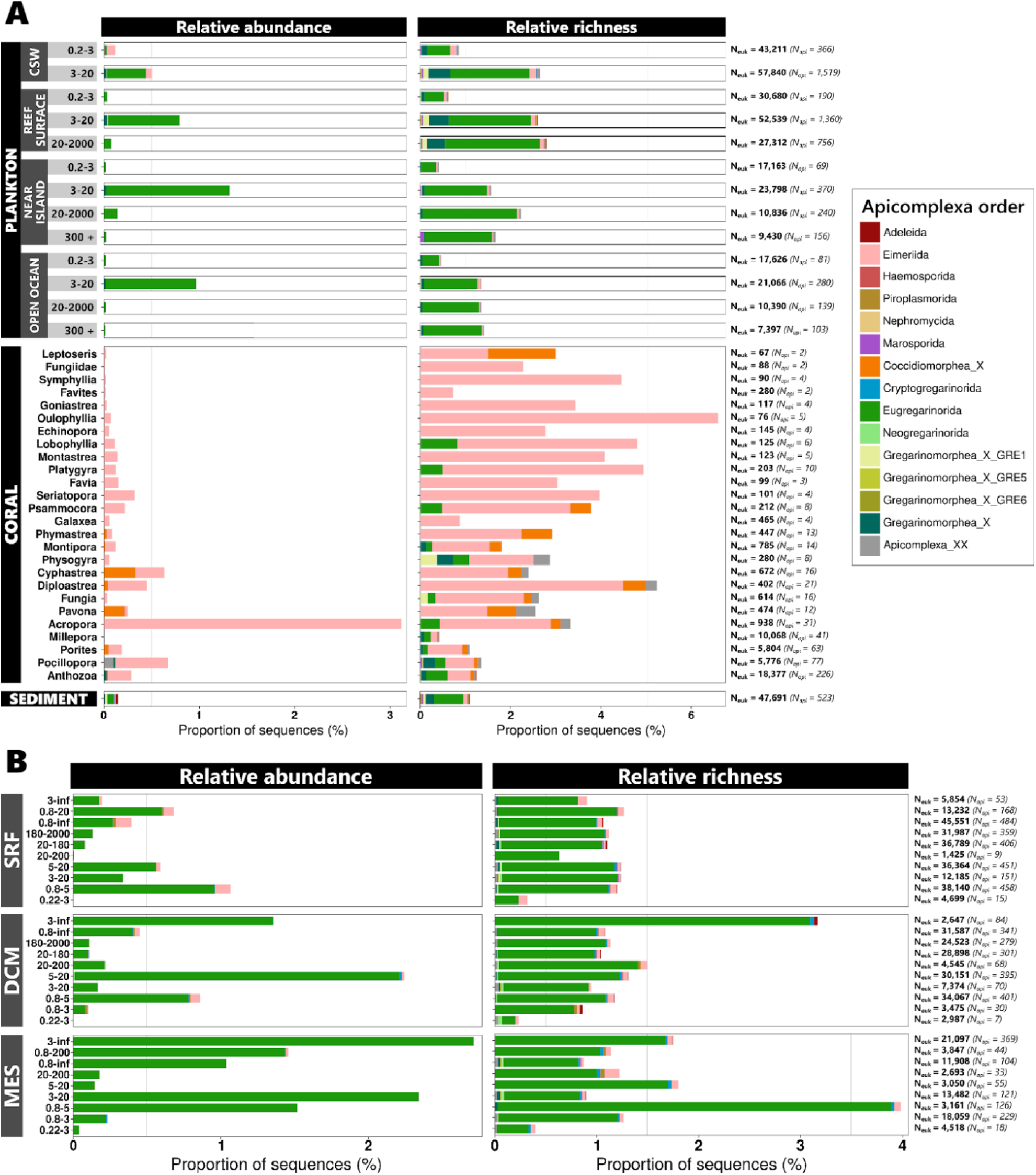
Relative abundance and richness of Apicomplexa across eukaryotic communities sampled during the Tara Pacific and Tara Oceans expeditions. (A) Tara Pacific dataset. Relative abundance (left) and relative richness (right) of Apicomplexa orders across different sample types, including coral hosts, plankton and sediment samples. Coral-associated samples are further subdivided by coral host genus, whereas plankton samples are structured according to their proximity to the reef and by the size fraction. CSW refers to coral surrounding water. (B) Tara Oceans dataset. Relative abundance (left) and richness (right) of Apicomplexa orders in plankton samples collected at different depths and size fractions, including surface waters (SRF), deep-chlorophyll maximum (DCM) and mesopelagic zone (MES). In both panels, relative abundance corresponds to the proportion of sequencing reads and relative richness corresponds to the proportion of ASVs. Colors represent Apicomplexa orders according to the PR2 taxonomy [51]. N(euk), number of eukaryote ASVs; N(api), number of apicomplexan ASVs.

**Supplementary Fig. 2.**
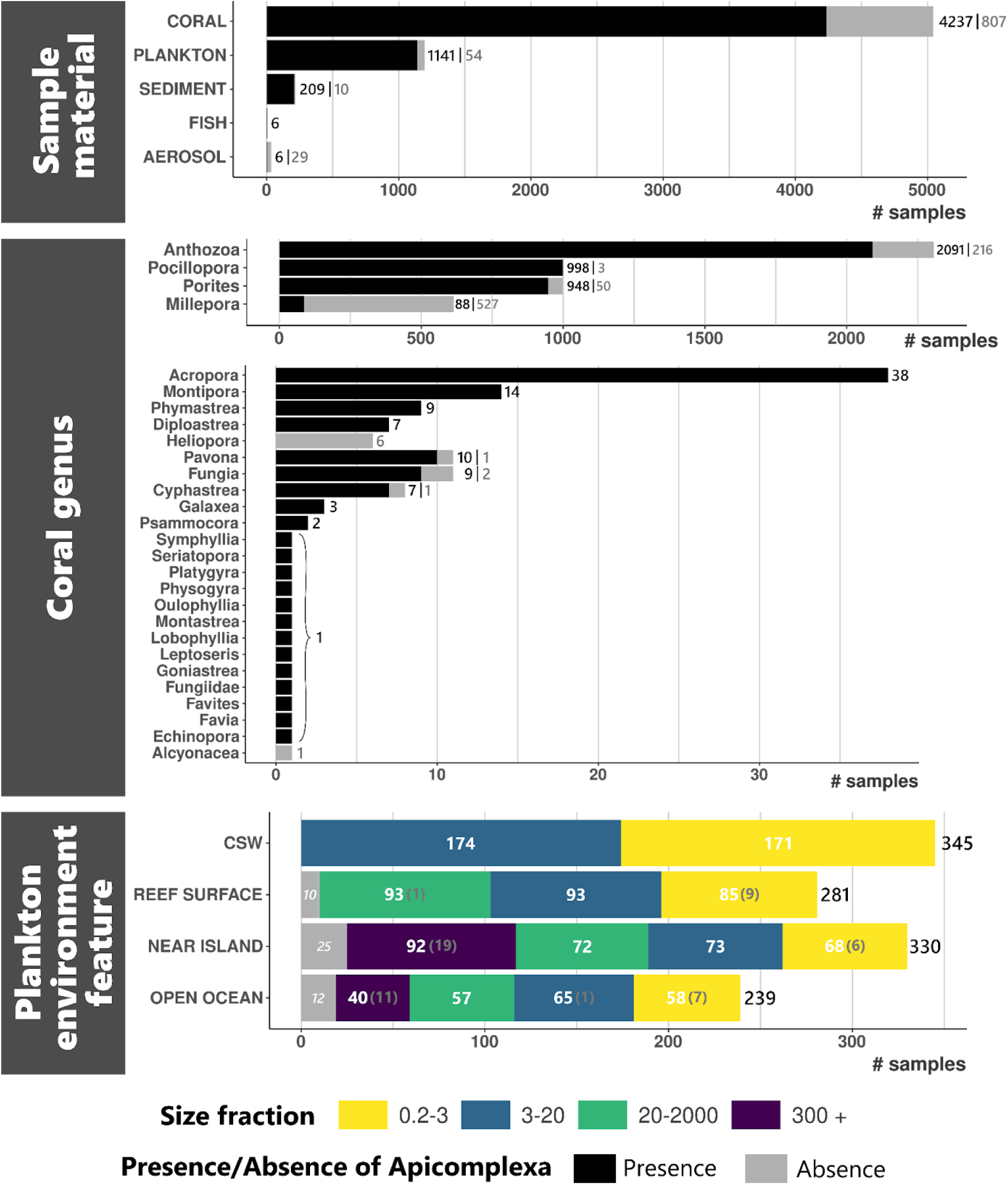
Distribution of samples according to ecological context and Apicomplexa detection. Horizontal barplots show the number of samples across sample types (top), coral host genera (middle), and planktonic environmental features and size fractions (bottom), separated by the presence (black) or absence (grey) of Apicomplexa. Numbers indicate absolute sample counts. In the coral panel, host genera are displayed on two x-axis scales to accommodate large differences in sampling effort among genera. In the plankton panel, stacked bars represent size fractions (colour-coded) allowing comparison of Apicomplexa detection across environmental contexts and particle size classes. “Anthozoa” stand for “unidentified Anthozoa”.

**Supplementary Fig. 3.**
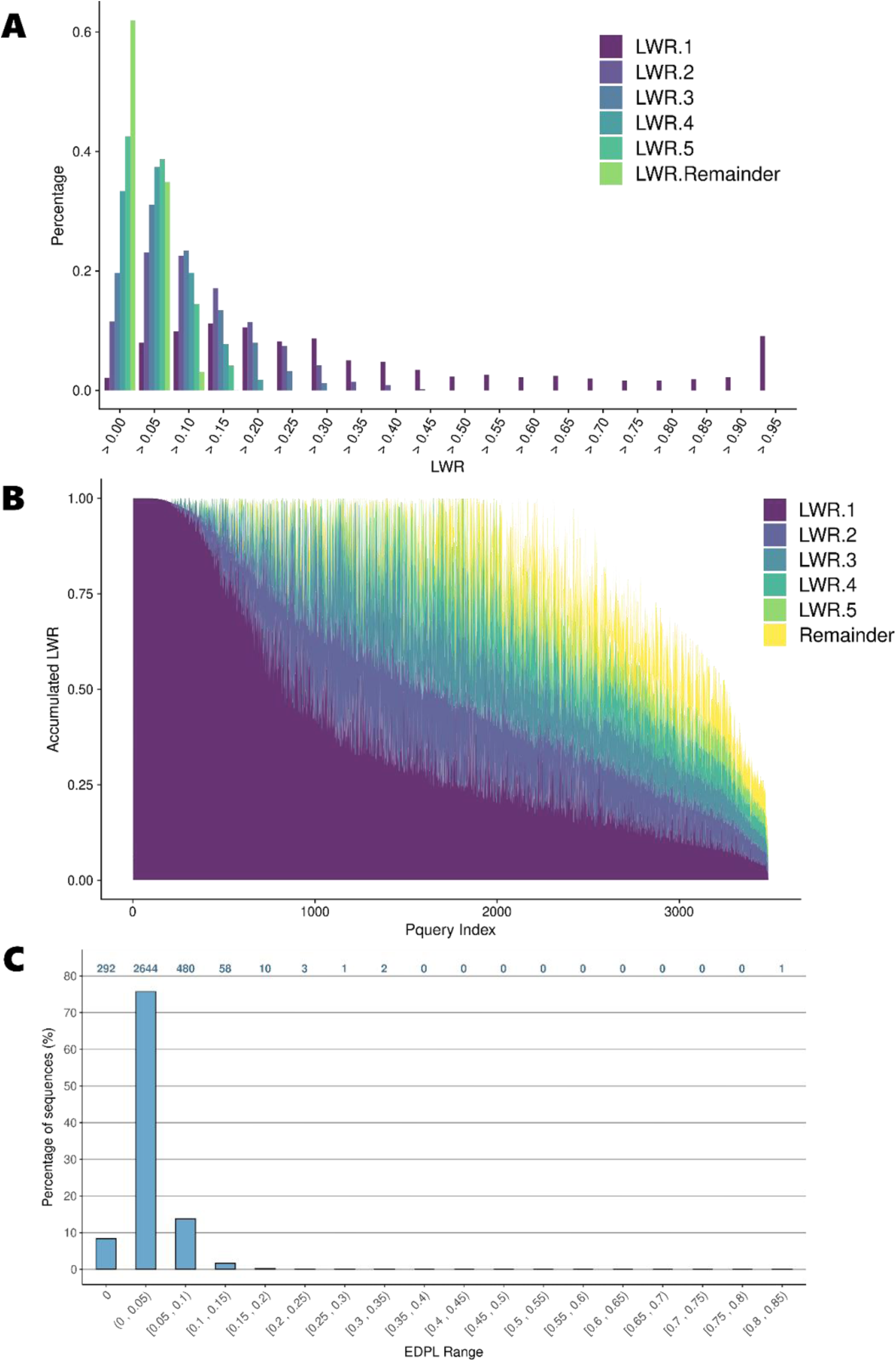
Phylogenetic placement support of Apicomplexa ASVs from Tara Pacific and Tara Oceans. Summary statistics of phylogenetic placement for Apicomplexa ASVs from Tara Oceans and Tara Pacific mapped onto the reference Apicomplexa tree. (A) Distribution of likelihood weight ratio (LWR) for the first five placement positions and the remaining placements. (B) Accumulated LWR across all query sequences, ordered by decreasing support of the best placement. (C) Distribution of expected distances between placement locations (EDPL), reflecting the spatial dispersion of alternative placements along the reference tree. The distribution of placement scores and the predominance of low EDPL values collectively indicate a satisfactory placement quality for most sequences globally, with around 25% of sequences exhibiting particularly robust support for their primary placement (LWR1 > 0.70).

**Supplementary Fig. 4.**
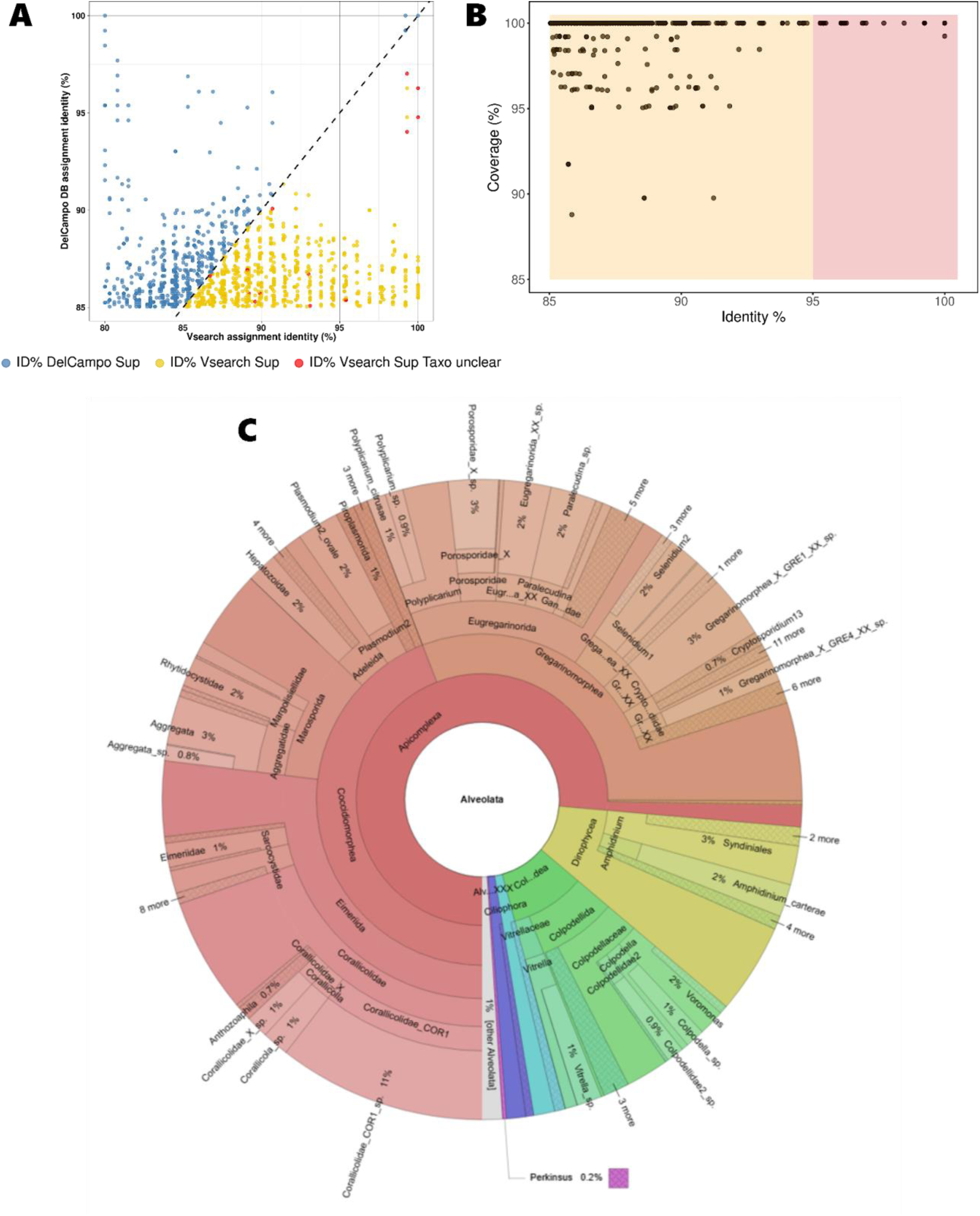
Identification and phylogenetic assignment of taxonomically unknown Tara Pacific ASVs. These ASVs were initially classified at high taxonomic ranks (above Apicomplexa) using IDTAXA but showed significant similarity to Apicomplexa reference sequences. (A) Comparison of sequence identity percentages between these Tara ASVs and Apicomplexa sequences either from the reference tree or from the PR2 database. Only best BLAST matches with >= 80% query coverage and >= 85% identity were retained. Points represent 2,081 BLAST matches corresponding to 1,329 unique ASVs. Colours indicate ASVs for which identity to the reference sequences is higher or equal to VSEARCH identity (blue), VSEARCH is higher but associated with an imprecise taxonomic assignment (red), or VSEARCH identity is higher with a clear assignment (yellow). (B) Identity-coverage relationship for the final subset of retained ASVs (715 ASVs; 1,071 BLAST matches); All retained sequences show high similarity to Apicomplexa reference (> 85% identity), with the majority exhibiting high alignment coverage (> 95%), supporting their classification as plausible Apicomplexa candidates. (C) Krona representation of placement-based taxonomic assignments for the 715 retained ASVs, inferred from phylogenetic placement onto the Apicomplexa reference tree. ASVs placed within the Apicomplexa clade are shown in red, whereas ASVs placed with reference outgroup lineages are shown in other colors.

**Supplementary Fig. 5.**
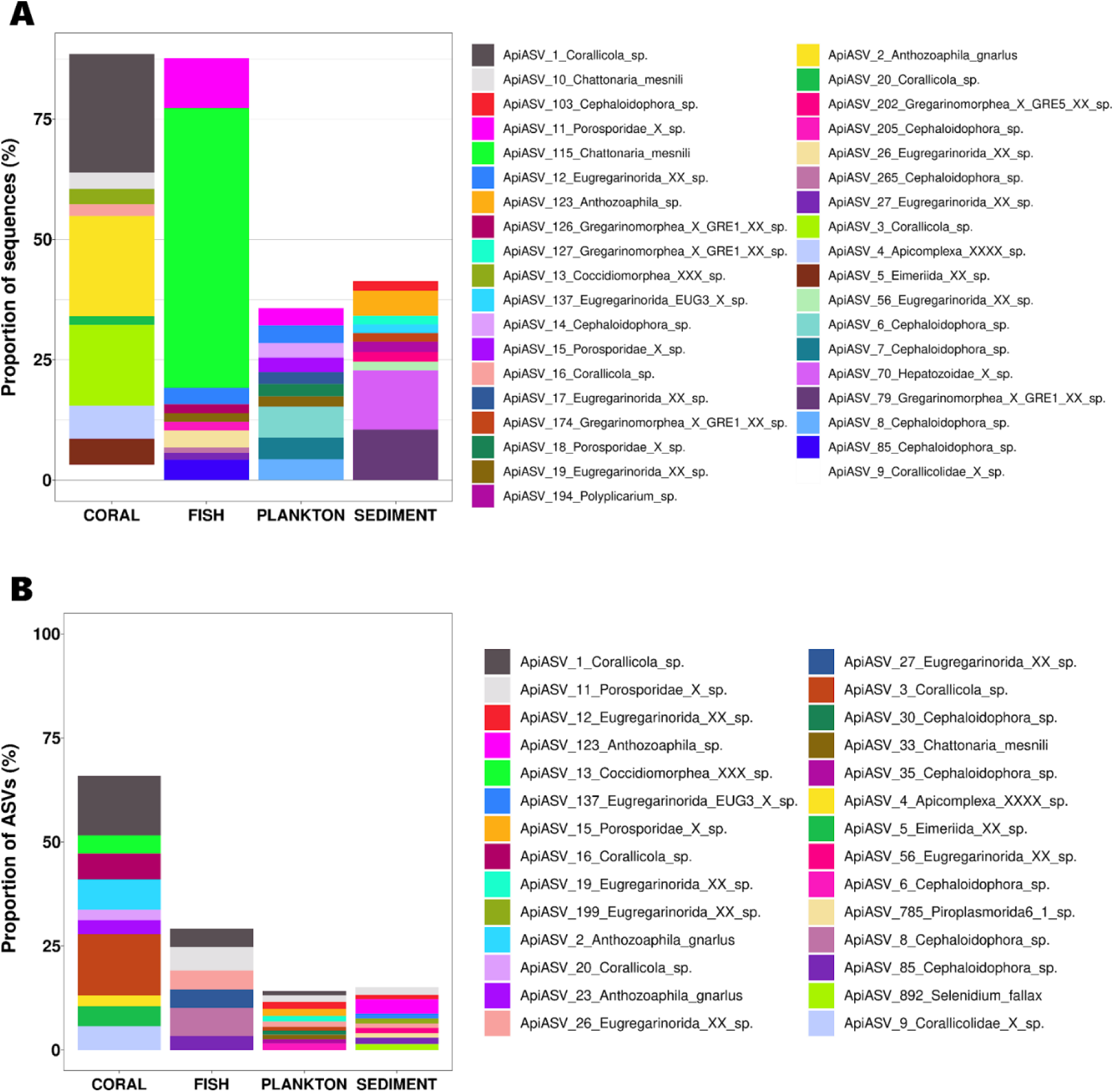
Relative abundance and richness of the ten most dominant Apicomplexa ASVs across environments. Stacked bar plots show the distribution of the ten most abundant Apicomplexa ASVs based on read abundance (A) and Apicomplexa lineages are the most diverse (in terms of ASV counts) the ten most ASV-rich Apicomplexa taxa (B) across environmental compartments (coral, fish, plankton and sediment). Bars represent the proportion of sequences (A) or the proportion of ASVs (B) attributed to each ASV lineage within each compartment.

**Supplementary Fig. 6.**
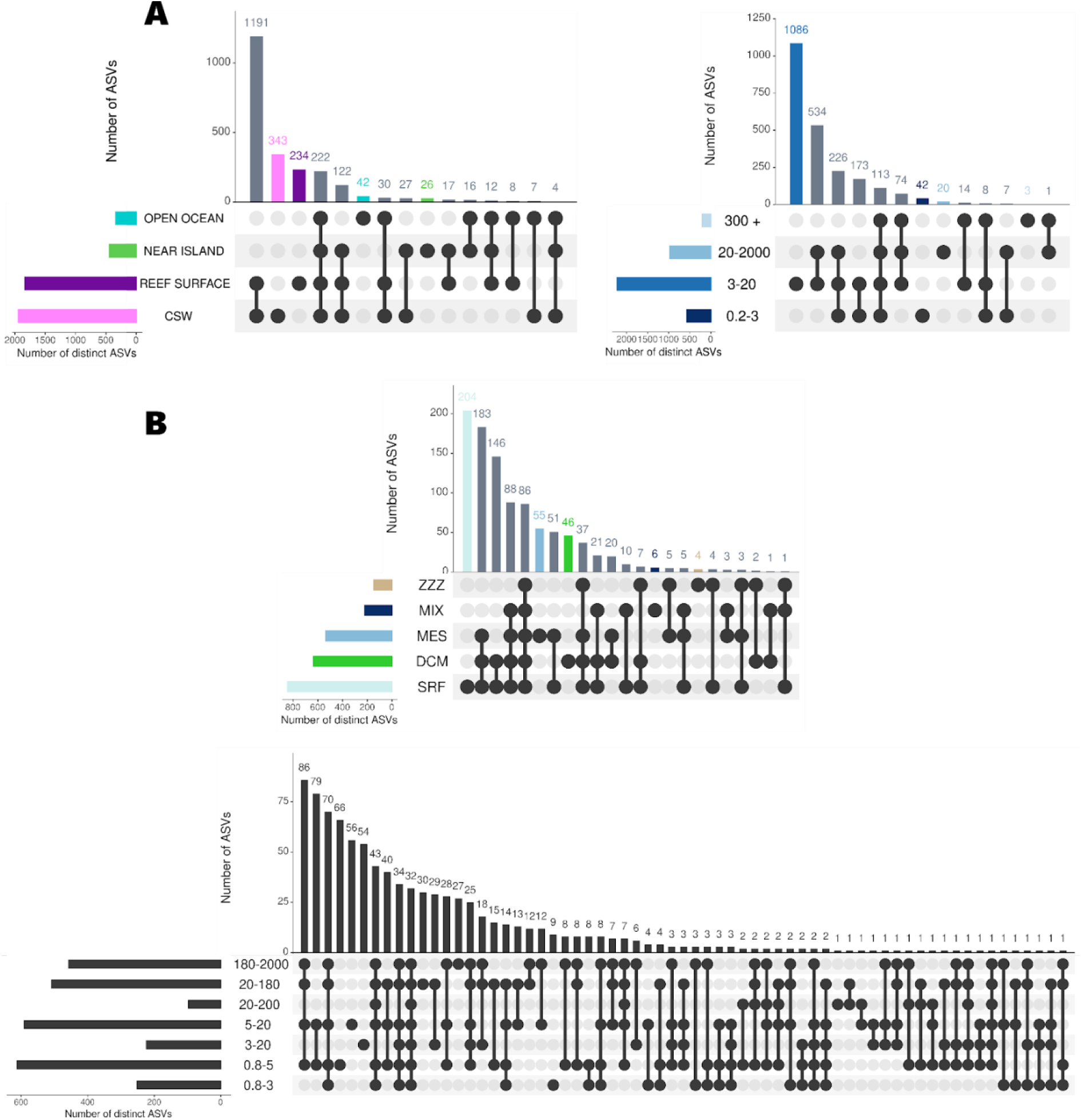
Shared and unique Apicomplexa ASVs across planktonic environments, size fractions and depth layers. UpSet plots show the number of shared and unique Apicomplexa ASVs across planktonic samples. (A) Tara Pacific planktonic samples, partitioned by oceanic regions (left) and size fractions (right). (B) Tara Oceans samples, partitioned by depth layers (top) and size fractions (bottom). Bars indicate the number of ASVs per intersection, with connected dots representing the corresponding combinations of categories. SRF, surface waters surface; DCM, deep chlorophyll maximum; MES, mesopelagic waters; MIX, surface mixed layer; ZZZ: undefined or non-standard sampling layer, used for samples that could not be assigned to a specific depth category.

**Supplementary Fig. 7.**
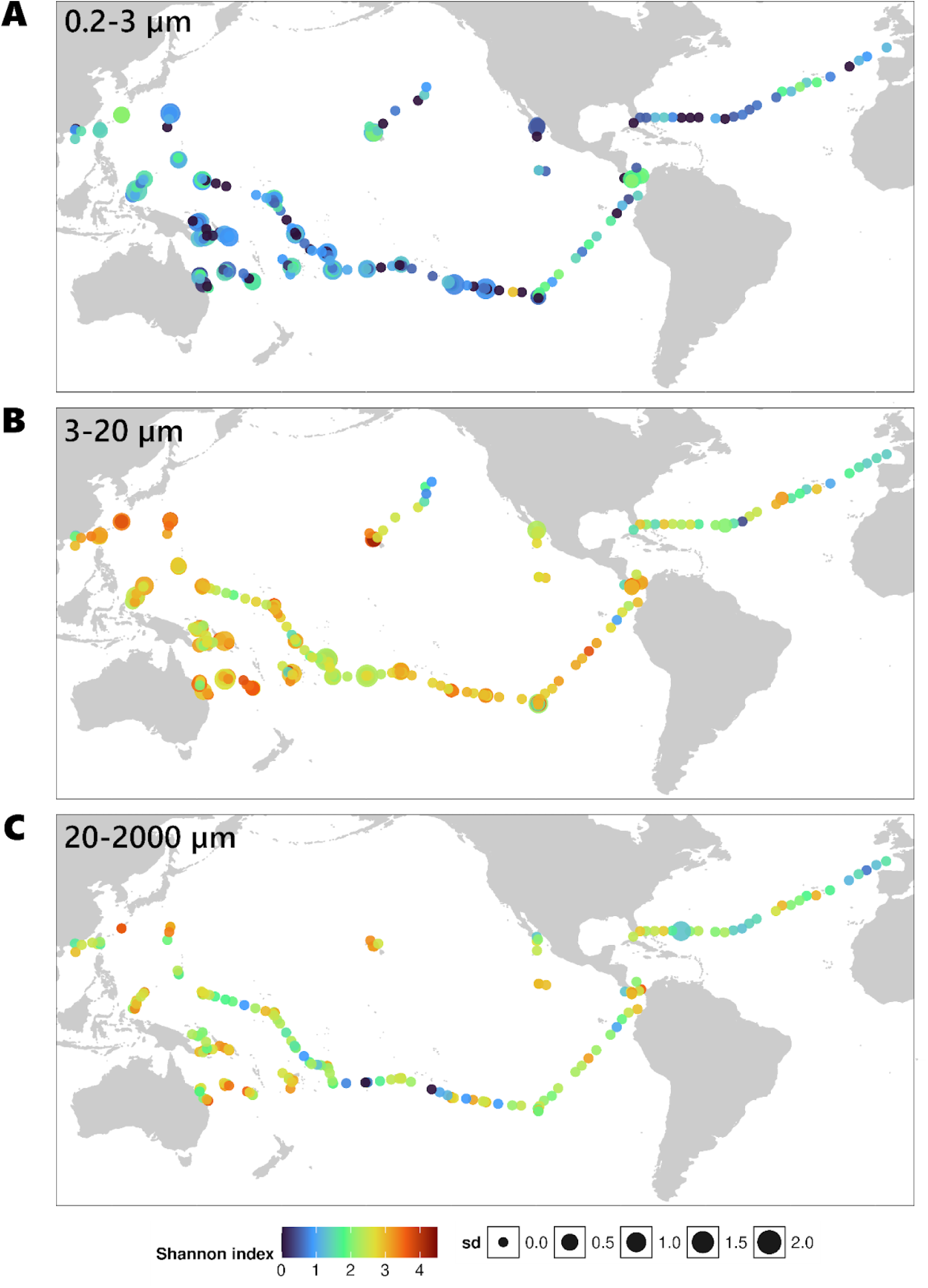
Global distribution of planktonic Apicomplexa diversity across size fractions. Global maps showing the spatial distribution of Shannon diversity indices for planktonic Apicomplexa across three size fractions: 0.2–3 µm (A), 3–20 µm (B) and 20–2000 µm (C). Each point represents the mean Shannon index calculated for a given geographic coordinate, integrating all planktonic samples collected at that location within the corresponding size fraction. Samples from different environmental contexts (open ocean, near island, island and coral surface waters) were pooled for visualization. Dot color indicates the mean Shannon diversity, while dot size reflects the standard deviation among samples sharing the same coordinates. Land masses are shown in grey.

**Supplementary Fig. 8.**
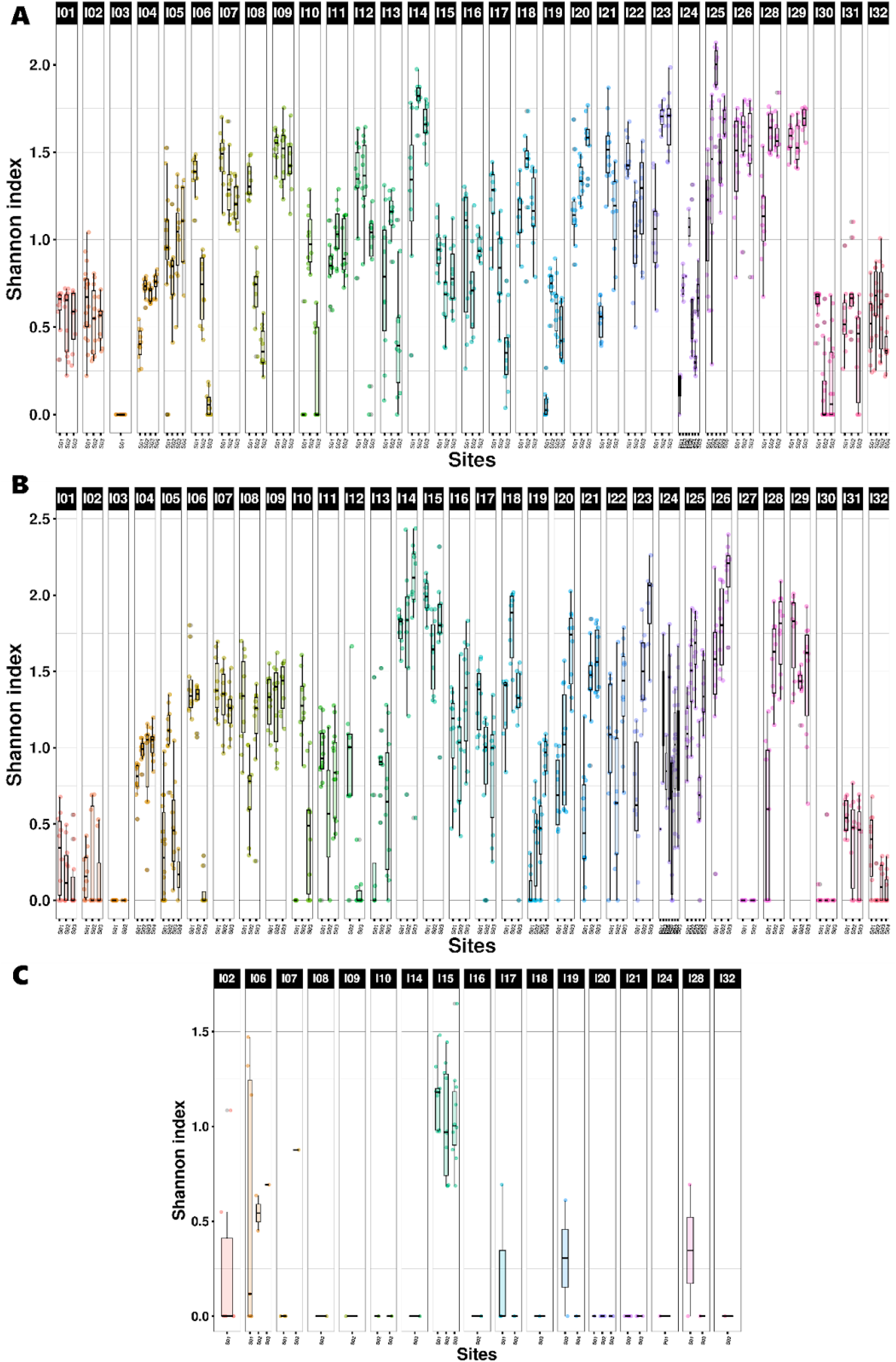
Distribution of Shannon diversity across sampling sites for coral-associated Apicomplexa. Boxplots show the distribution of Shannon diversity index values across sampling sites, grouped by island, for the coral genera *Pocillopora* (A), *Porites* (B) and *Millepora* (C). Points represent individual samples.

